# A geometric criterion links HIV-1 capsid topography to its biophysical properties and function

**DOI:** 10.64898/2025.12.01.691629

**Authors:** Wenhan Li, Craig A. Peeples, Juan S. Rey, Juan R. Perilla, Reidun Twarock

## Abstract

Mathematical models of virus capsid structure are pillars of modern virology, aiding the understanding of viral mechanisms and the design of antiviral interventions. Traditionally, the HIV-1 capsid core geometry is represented as a fullerene lattice, akin to the icosahedral models of spherical viruses in Caspar-Klug theory. However, recent studies revealed that many viral capsids deviate from such idealised lattices, with important functional implication. Here we show that this is the case also for the conical HIV-1 core geometries, in which the hexamer and pentamer boundaries form a pseudo-tiling rather than a perfectly aligned fullerene network. We introduce a triangular geometric criterion that quantifies local deviations of an HIV-1 atomic model from its idealised fullerene backbone. Using this criterion, we present that this difference in geometric organisation between idealised (fullerene) and actual (data-derived) capsid model has implications for the capsid’s biophysical properties. We also discuss the use of the geometric criterion as a predictive tool regarding cofactor binding and implied geometric changes in the capsid surface coupled to the interfacial frustration response. Our results establish a quantitative framework linking capsid geometry, curvature, and biophysical function, offering new perspectives for assembly inhibitor design and lentiviral vector engineering.

## Introduction

The human immunodeficiency virus type 1 (HIV-1) is the predominant cause of the global HIV epidemic and acquired immunodeficiency syndrome (AIDS)^1,2^. During viral infection and replication, its protein shell, or capsid core, performs essential functions including encapsulating and shielding the viral genome from degradation, serving as a reaction vessel for reverse transcription, and facilitating its trafficking and nuclear import^3–5^. Due to these important roles, the capsid has become a central target for antiviral drug design^4–6^. On the other hand, HIV-based lentiviral vectors are also exploited as delivery systems for gene therapy applications, to treat genetic diseases, primary immunodeficiencies and cancers^7–9^. A better understanding of the biophysical properties and (dis)assembly mechanisms of the capsid core is therefore crucial for both antiviral and therapeutic applications.

As structure and function are intertwined, the mathematical modelling of viral capsid structure plays a significant role in virology. The spherical capsid models from Caspar-Klug theory are a primary tool to classify and understand virus structure^10,11^. However, recent analyses show that many spherical viral shells deviate from these idealised lattices, with important functional consequences^12^. For example, different viruses with the same number of capsid proteins, that would traditionally be represented by the same hexagonal lattice architecture in Caspar-Klug theory, can have very different biophysical properties in terms of their resilience to fragmentation depending on their actual surface lattice type and relative orientation of their capsid proteins^13^.

Unlike the abundant spherical viruses exhibiting icosahedral symmetry, the HIV-1 capsid has a distinct conical morphology (Fig. 1a) composed of approximately 200 hexamers (shown in grey) and 12 pentamers (orange brown). It has traditionally been represented over the past decades as an idealised fullerene or hexameric lattice with pentameric insertions^14–17^ (Fig. 1b, d, e). Yet, local rotations and positional adjustments of hexamers and pentamers – arising from the formation of the dimer interfaces (red lines between capsomers in Fig. 1f-i) – result in imperfect alignment between adjacent capsomers (Fig. 1c, f-i). Such geometric mismatches suggest that an alternative model^12,13^ may be required to better capture its surface architecture. To examine this, we compare three geometric frameworks: the fullerene lattice (a regular hexagonal network with 12 pentagonal insertions); the Kagome lattice (a rotated hexagonal and pentagonal arrangement forming interlaced triangles); and a data-derived lattice constructed directly using the atomic positions of an identical residue (as explained below) to reproduce the actual orientations of the hexameric and pentameric lattice units. In contrast to the fullerene lattice, the data-derived lattice explicitly incorporates the geometric mismatches between neighbouring capsomers. In this work, we show that this difference in geometric representation has consequences for how well the geometric model captures the biophysical properties of the HIV-1 capsid, such as those quantified via molecular frustration computations. This implies that choosing the correct geometric representation is paramount to best represent capsid functionality.

**Fig. 1.**
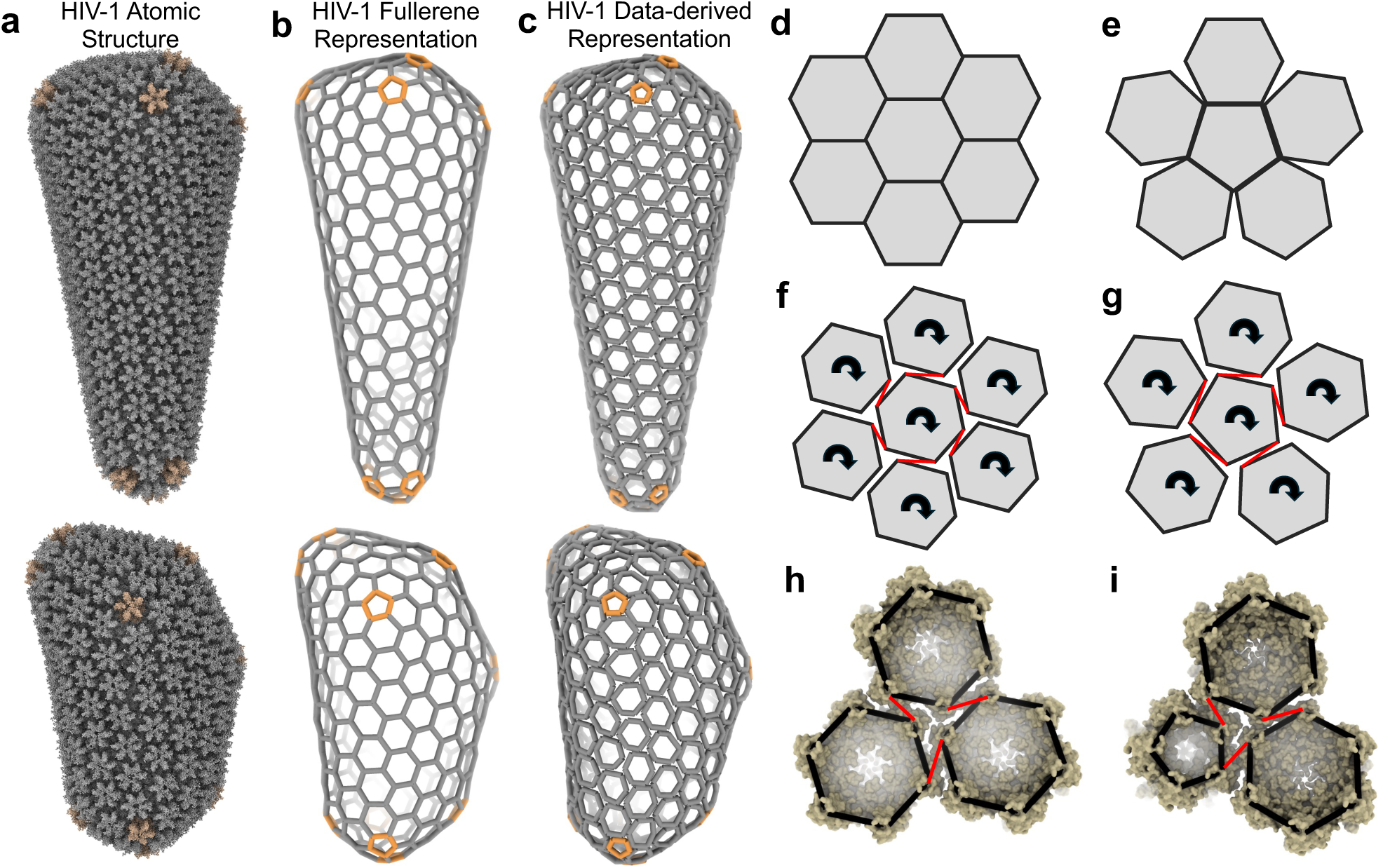
Comparison of an idealised (fullerene-style) representation with the capsid geometry of two HIV-1 cores. **a**, Experimentally derived atomic models of the HIV-1 capsid (vlp23 (top) and PDB: 3J3Y (bottom)). **b**, A fullerene-style representation of capsid geometry. **c**, Capsid geometry derived from the atomic positions of the CA hexamers and pentamers. CA hexamers and their respective geometric representations are coloured in grey while pentamers are coloured in orange. **d**-**g**, Schematic representation of the difference in geometric organisation between the fullerene-style (**b,d,e)** and data-derived (**c,f,g)** lattices around a hexagonal (**d**,**f**) and a pentagonal (**e**,**g**) unit, illustrating geometric mismatches between neighbouring capsomers in the data-derived lattice due to lattice gyration (**f**,**g**). **h**-**i**, Capsomers are shown superimposed on the data-derived lattice architecture in an inside-out view, with red lines indicating dimer interfaces.

Advances in computational biology and cryo-electron microscopy (cryo-EM) have enabled atomistic reconstructions of entire HIV-1 capsids following different modelling routines^16,18^. Such atomistic models are essential steps, in combination with experiment, in enabling a better understanding of the mechanisms underpinning viral life cycles^19–21^. In addition, computer simulations identify capsid assembly and maturation pathways in agreement with *in vitro* maturation data^22–24^, revealing the formation of a curled sheet in the bulk of the capsid as an intermediate step. Seam lines in the lattice architecture of the mature cone may originate from the junctions of these curled sheets. This suggests that local curvature and lattice geometry play a pivotal role in capsid function. However, the quantitative characterisation of CA lattice geometry – and its impact on the function of the conical HIV-1 capsid – remains underexplored. A previous study introduced three orientations of hexamers with high, intermediate, and low curvature in a tubular surface lattice^25^. Yet, describing curvature within the asymmetric conical lattice is considerably more complex. Although areas surrounding pentamers display higher curvature as expected, the overall lattice lacks simple geometric regularity: adjacent capsomers exhibit tilting and twisting relative to one another^26^, and individual capsomers possess intrinsic curvature themselves^27^.

Here, we introduce a local geometric criterion that quantitatively characterises the arrangement and deformation of the conical HIV-1 surface lattice. We first present that the geometric properties of different computationally generated carbon-fullerene cone structures, as captured by the geometric criterion, correlate strongly with their energy landscapes. We then extend this framework to the HIV-1 atomic model, linking geometric and biophysical properties of the capsid. Our analysis reveals that only the data-derived lattice model accurately reproduces the potential seam line location and associated molecular frustration patterns observed experimentally, in support of the hypothesis that the geometric representation influences the inferred biophysical landscape of the capsid. By computing molecular frustration^28,29^ at dimer interfaces, we show that regions of high frustration align with the seam line exclusively in the data-derived lattice, highlighting the importance of the choice of lattice representation in a coarse-grained model. We also show that the geometric criterion distinguishes between capsids reconstructed via different protocols, indicating that assembly history may imprint characteristic geometric signatures on the capsid lattice, that can be quantified by the geometric index introduced here.

To illustrate the predictive power of our geometric index, we extend our analysis to host cofactor and small molecule binding. Cofactors such as cytoplasmic cyclophilin A (CypA)^30,31^, and small molecular weight compounds like lenacapavir (LEN)^32–34^ interact with specific surface sites of the CA lattice. It has been suggested that such binding sites may depend on local geometric features of the surface lattice^27^, but this association has not yet been made precise. Here we show that the geometric criterion correlates with local curvature and captures the site-specific geometric and energetic responses to ligand binding at the two primary binding sites^35–37^. Binding at the CypA site induces a local curvature increase accompanied by elevated frustration values at nearby dimer interfaces, whereas binding at the phenylalanine-glycine (FG)-motif pocket produces lattice flattening and reduced interfacial frustration values. These contrasting behaviours reveal a direct link between local geometry, curvature modulation, and cofactor function, suggesting that HIV-1 exploits surface topography to regulate capsid mechanics and structural dynamics during its life cycle.

## Results

### The geometric index of hexagonal surface lattice architectures

To quantify the geometric organisation of a hexagonal surface lattice architecture such as a fullerene cone, we introduce a triangular geometric index that captures local topographic features. As illustrated in Fig. 2a, a characteristic triangle (purple dashed lines) is initially defined at any threefold axis surrounded by three regular hexagons. Because hexagons in fullerene cones are typically distorted as they are part of a surface lattice, we also constructed analogous characteristic triangles at each local threefold axis of a fullerene cone (Fig. 2b & 2c). For each triangle, the shortest edge is highlighted in green. Its length and orientation in the surface lattice indicate the directions of maximum lattice compression. These shortest edges tend to align in the surface lattice, revealing local patches where hexagons are deformed collectively in a consistent direction to form smooth sheets. We therefore refer to the shortest edge at each local threefold axis as the geometric index, and its definition below will therefore capture both the length and the orientation of the shortest edge.

**Fig. 2.**
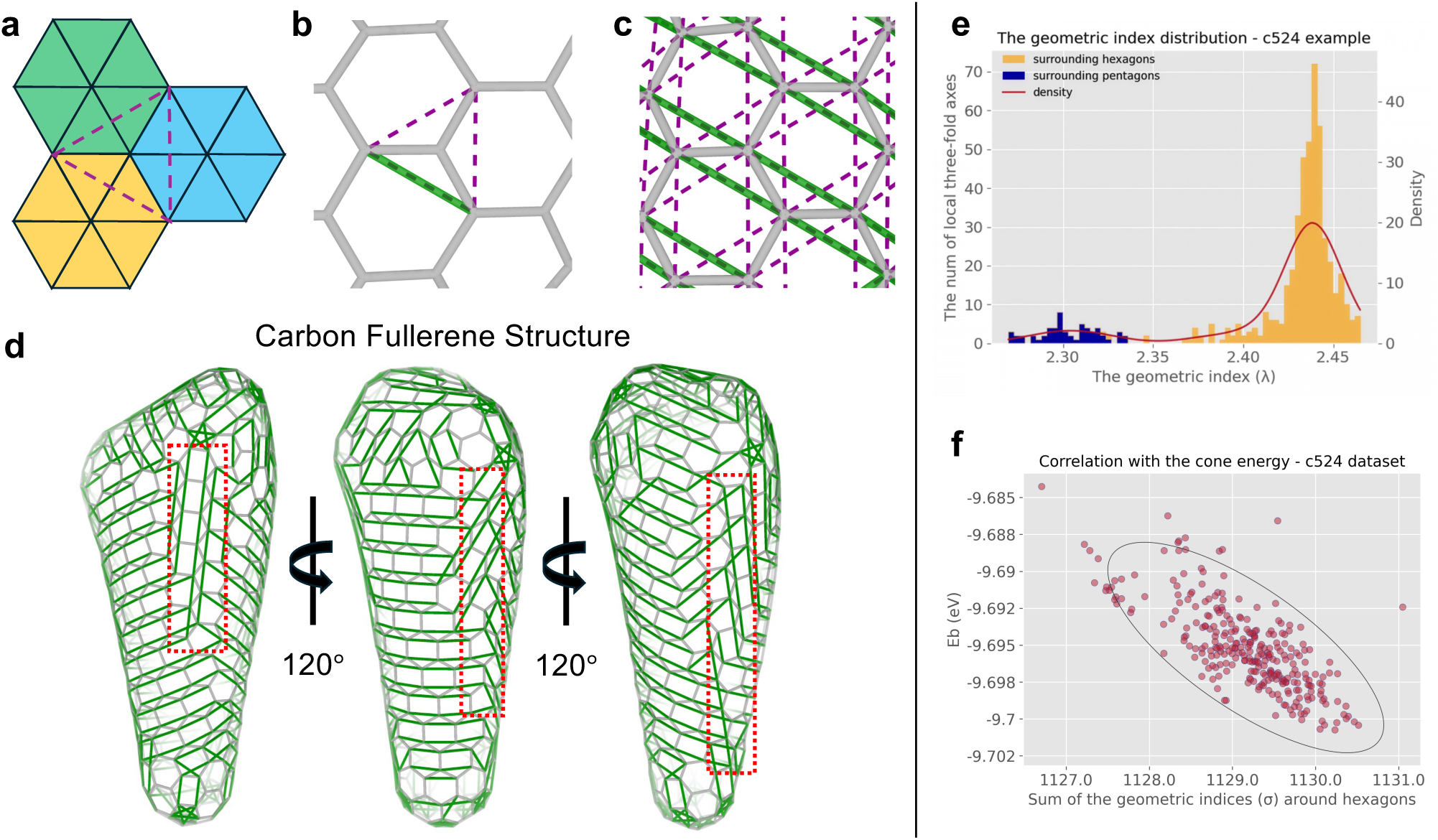
The characteristic triangle and the geometric index applied to the carbon fullerene-lattice representation. **a**, A characteristic triangle (purple dashed lines) at a threefold axis surrounded by three regular hexagons. **b**, A characteristic triangle at a local threefold axis of a fullerene lattice (light grey solid lines), where the shortest edge (i.e., the geometric index) is highlighted in green. **c**, The characteristic triangle and its geometric index are extended to each local threefold axis of a fullerene lattice. **d**, The geometric indices (green lines) on the surface of a fullerene cone (C_524_ dataset). The geometric indices partially align, forming local sheets. These sheets are joint together at seams (highlighted by red dashed frames). **e**, The distribution of the geometric indices across the cone in **d**. **f**, The sum of the geometric indices surrounding hexagons correlates with cone energy (shown here for cones in the C_524_ dataset). Source data are provided as a Source Data file.

For a fullerene cone model containing *m* vertices, representing the positions of the carbon atoms, each local threefold axis is labelled by *i* ∈ *I* = {1, 2, …, *m*}. The magnitude of the geometric index at the *i*^th^ local threefold axis is then defined as

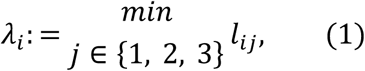

where *l*_ij_ denotes the length of the *j*^th^ edge of the characteristic triangle. The orientation of the geometric index is defined as a bidirectional unit vector *v*^↔^_i_pointing along the corresponding shortest edge. The complete set of geometric indices is therefore

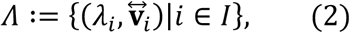

in which each element encodes both the local amount (λ_i_) and the direction (*v*^↔^_i_) of lattice deformation.

Building on prior work^16^ where fullerene models serve as structural backbones for HIV-1 capsid reconstruction, we generated carbon-based fullerene cones using the dedicated Fullerene program^38–41^, followed by quantum tight-binding density functional theory (TB-DFT) energy optimisation and capsid structure determination (see Methods). The Fullerene program was used to generate fullerene models by adjusting ring spiral pentagon indices (RSPIs) in small increments. Modified input files were tested for validity based on carbon count and RSPI rules, with parallel searches identifying a small set of valid candidates. Three datasets of cone fullerene structures were constructed with 352 (C_352_), 452 (C_452_), and 524 (C_524_) carbon vertices, respectively. Structural polymorphism within each dataset arises from variations in the positions of pentagons at both ends of the cone.

To examine how the geometric organisation of fullerene structures relates to their intrinsic energy, we mapped the geometric indices onto the surfaces of the computational models. Representative examples from C_524_ and C_352_ are shown in Fig. 2d and Supplementary Fig. 1a, respectively. Green lines denote geometric index directions, which partially align to form local sheets separated by seam lines. Typically, two to three seams appear in the bulk of each cone, joining adjacent sheets (red dashed frames). In the computationally generated carbon fullerene cones, these multiple seams arise because the fullerene construction algorithm explores admissible geometric configurations without imposing biological or chemical constraints on assembly. By contrast, the HIV-1 capsids discussed below rely on the atomic positions of the capsid proteins and MD simulations. In the fullerene cone models, seams can therefore be interpreted as boundaries between locally aligned sheet-like regions that minimise distortion within each region while accommodating global curvature. In the viral capsids we observe fewer seams, consistent with previous simulations^24^. This is likely the case as protein interactions bias the system towards configurations with fewer seams, given the energetic cost associated with seam formation.

The distribution of the scalar component of the geometric index (its length) is bimodal in both examples (Fig. 2e and Supplementary Fig. 1b). The indices computed at threefold axes surrounding pentagons (blue histograms in Fig. 2e and Supplementary Fig. 1b) are typically smaller than those around hexagons, indicating that the diagonals of the pentagons are generally shorter than those of the hexagons. Notably, the sum of the geometric indices σ = ∑_i∈*I*_ λ_i_ for each cone is strongly correlated with the intrinsic energy of the structure *E*_b_ (Fig. 2f and Supplementary Fig. 1c), highlighting the role of the local geometric organisation in determining the overall energetic landscape of the fullerene cones.

### Distinct topographies revealed by the geometric indices across HIV-1 capsid models

To compare the geometric organisations of different HIV-1 capsid models, we constructed three geometric representations based on the capsid atomic model vlp23^18^: a fullerene, a Kagome, and a data-derived surface lattice with vertices derived from the atomic positions of an identical residue in the dataset (see Methods). The basic lattice building blocks are shown in Fig. 3a-c (left), where black dots represent the vertices of the fullerene lattice, and grey dots those of the Kagome lattice. Coloured hexagons of equal size indicate the orientations of the hexagonal units in each lattice, with characteristic triangles outlined by purple dashed lines. The data-derived lattice can be considered an intermediate structure between the fullerene and Kagome model: the hexagons and pentagons in the Kagome lattice are rotated by ∼30° relative to those in the fullerene lattice, while those in the data-derived lattice (Fig. 3c & 3d) exhibit an intermediate rotation – less than 30° – positioning them geometrically between the two idealised lattices.

**Fig. 3.**
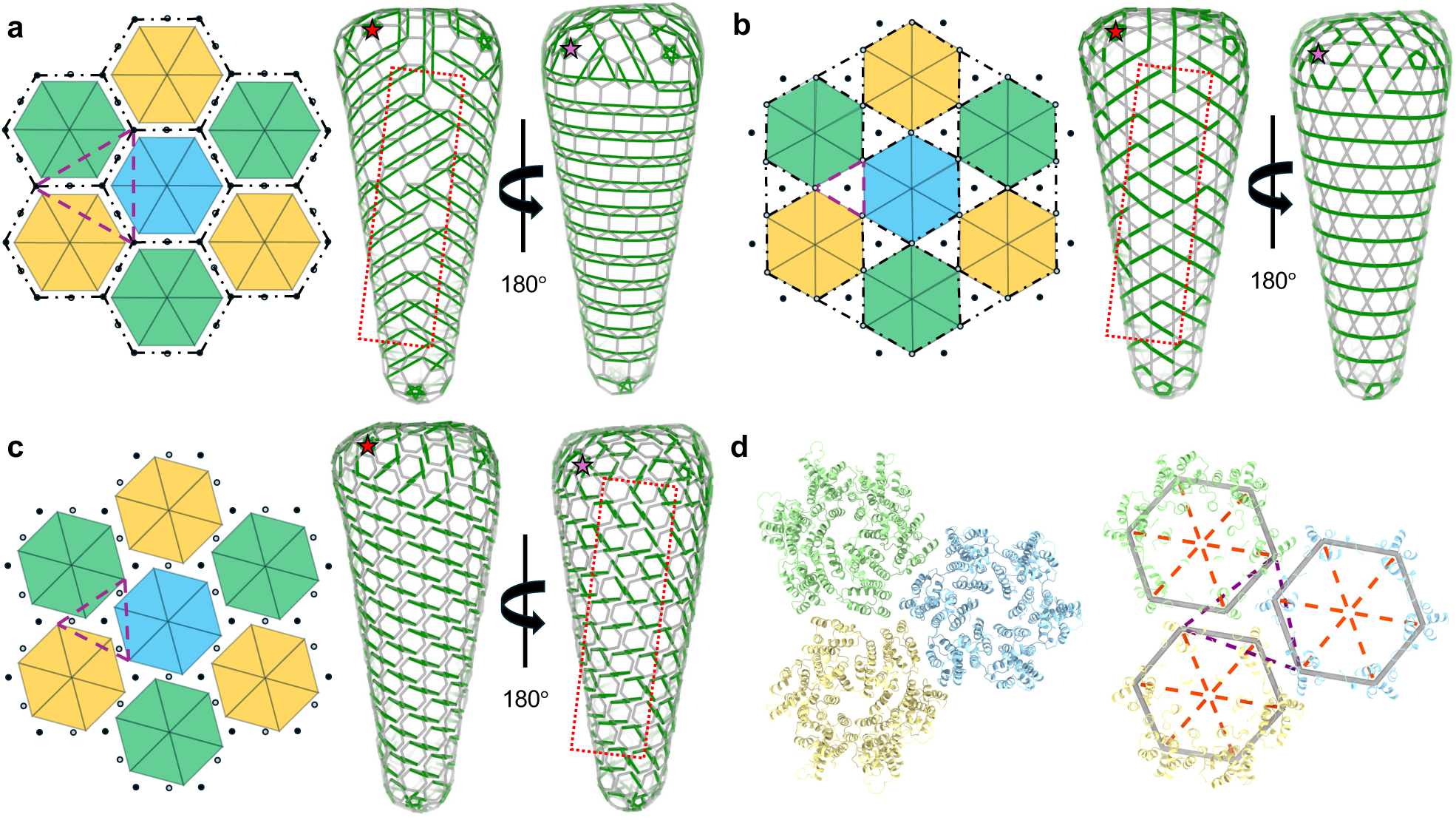
The distinct topographies of different HIV-1 capsid models, revealed by their geometric indices, in comparison. **a**, Left: The characteristic triangle of a fullerene-style lattice (purple dashed lines). Right: the reconstructed fullerene lattice for vlp23 (see also Supplementary Fig. 3 & 5). A seam line in the bulk is indicated by a red dashed box. A pentamer is labelled by a red star for reference to the other cases. **b**, Left: The characteristic triangle for a Kagome lattice model, where each hexagon is rotated by 30° about the centre of the original hexagon in **a**. The characteristic triangles (purple dashed lines) correspond to the triangular gaps produced by such rotations between three mutually adjacent hexamers. Right: the reconstructed Kagome lattice for vlp23 (see Supplementary Fig. 3 & 5). A seam line, located on the same side as in **a**, is indicated by a dashed box. **c**, Left: a pseudo-tiling mimicking the actual organisation of the CA hexamers; it is a geometry in between a fullerene and Kagome lattice, with hexagons rotated by less than 30°. The characteristic triangles (purple dashed lines) are of intermediate size compared with the other scenarios, with vertices situated between the fullerene (black dots) and Kagome lattice (grey dots). Right: the corresponding lattice geometry; the seam line in the bulk is on the opposite site compared with the other cases as can be seen with reference to the pentamer labelled by a light purple star. **d**, Left: three hexamers shown in different colours. Right: the profile of these three hexamers (light grey hexagons), constructed using N195 C_α_. Orange dashed lines correspond to the diagonals, and the purple dashed lines indicate the characteristic triangle.

Side views of the three capsid models are shown in Fig. 3a-c (right), where the red and the purple stars mark the positions of pentamers on opposite sides of each structure. The geometric index orientations *v*^↔^, represented by green lines, delineate the surface topography of each model. As indicated by the red dashed boxes, the seam lines identified by the geometric indices occur on the same side of the fullerene and Kagome surface lattices (red star) but are located on the opposite side in the data-derived model (purple star). This shows that distinct geometric models produce markedly different surface topographies, even when modelling the same capsid, and the geometric index provides a robust descriptor for capturing such differences. In subsequent sections, we correlate the topography encoded by the geometric index with the capsid’s biophysical properties in support of the hypothesis that the data-derived model is a more accurate geometric representation.

### Correlation between capsid topography and molecular frustration at the dimer interfaces

To probe the local biophysical properties of the HIV-1 capsid, we analysed the molecular frustration patterns at the dimer interfaces by computing the configurational frustration index for all contacts involving the H9 helices using the Protein Frustratometer framework^28,29^. The configurational frustration index was chosen because it better reflects protein-protein interface energetics^42,43^. Briefly, the frustration of a contact between residues *i* and *j* is defined based on the energy difference between the native interaction and a set of decoy interactions, expressed as a Z-score. Highly frustrated contacts correspond to unfavourable native interactions relative to the decoys (represented by smaller Z-scores), whereas minimally frustrated contacts correspond to favourable ones (represented by larger Z-scores). High local frustration often coincides with regions critical for functional binding^42,43^ or allosteric transitions^44^. Here, we defined the local frustration state γ_i_at the *i*^th^ dimer interface as the average frustration index of all contacts involving the H9 helices at that interface (see Methods).

Fig. 4a (Movie 1) illustrates the reference frustration pattern as a baseline for a representative CA dimer interface (PDB: 2KOD). Interactions involving H9 helices are shown as solid lines for direct contacts and dashed lines for water-mediated contacts, coloured red for highly frustrated, green for minimally frustrated, and grey for neutral bonds. Representative interfaces of hexamers in vlp23 corresponding to the lowest and highest local frustration states are shown in Figs. 4b & c (Movie 2 and 3), respectively. Highly frustrated interfaces exhibit a larger separation between the H9 helices, fewer interfacial contacts, and a higher proportion of water-mediated interactions, whereas low-frustration interfaces display tighter helix packing with more contacts. These relationships are quantified by the inverse correlation between local frustration and contact number (Fig. 4f) and by the positive correlation between frustration and the fraction of water-mediated contacts (Supplementary Fig. 7c). Consistently, interfaces with more contacts contain fewer highly frustrated and more minimally frustrated interactions (Supplementary Fig. 7d-f).

**Fig. 4.**
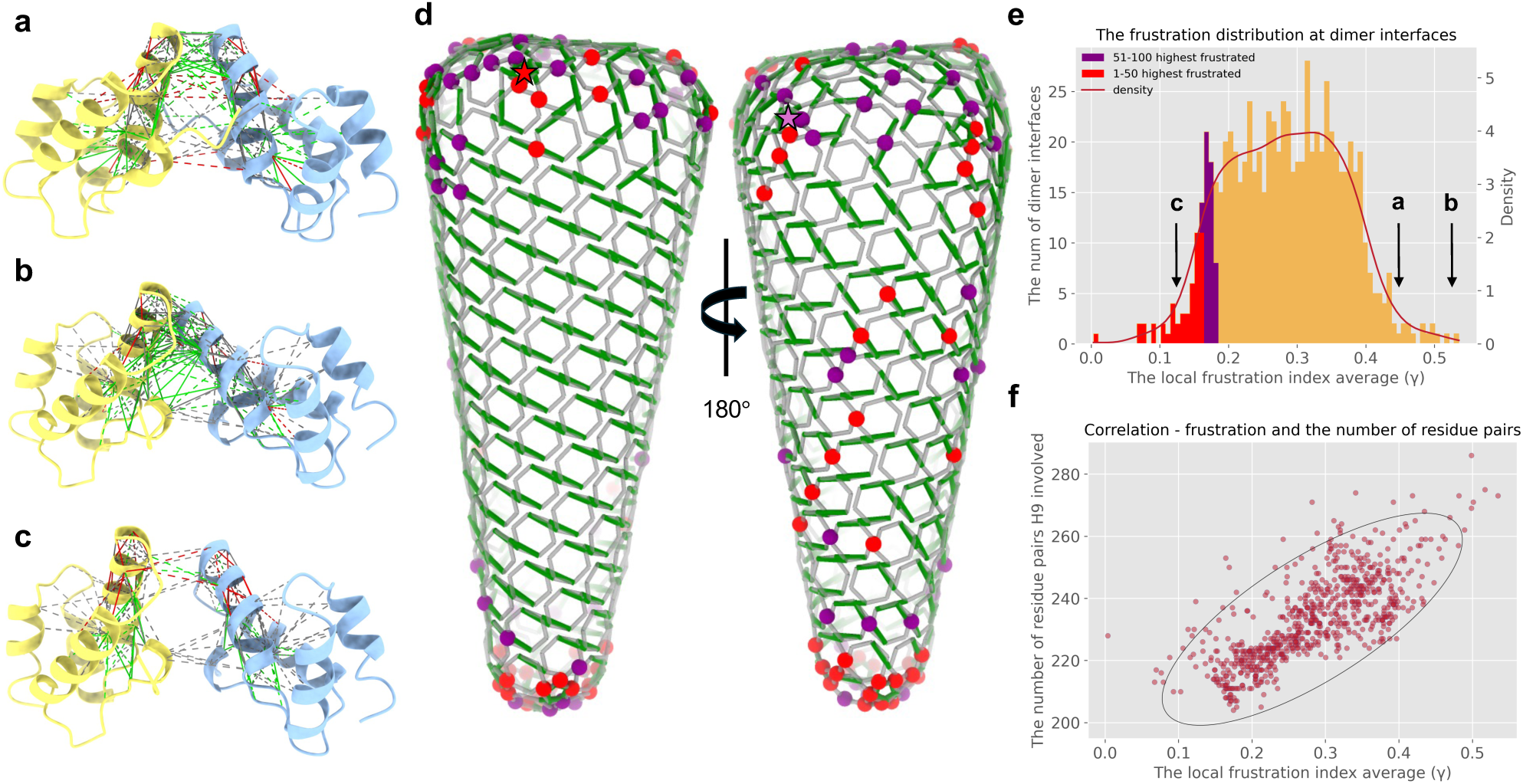
Molecular frustration at the dimer interface correlates with the capsid’s topography as encoded by the geometric index. **a**, The reference dimer interface (PDB: 2KOD); residue pairs involved in H9 helix interactions are shown as solid lines (direct contacts) or dashed lines (water-mediated contacts), coloured red for highly frustrated, green for minimally frustrated, and grey for neutral bonds. **b**-**c**, Representative dimer interfaces in vlp23 across two hexamers, showing the lowest (**b**) and highest (**c**) local frustration states. **d**, The data-derived geometric model of vlp23 with the geometric indices superimposed in green. Spheres indicate highly frustrated interfaces as follows: top 50 in red, and next 50 in purple. Side view showing the side with the pentamer marked by a red star (left), and the opposite side with the pentamer marked by a light purple star (right), corresponding to the seam side. The highly frustrated interfaces in the bulk are all located on the seam side. **e**, The distribution of the local frustration states across different dimer interfaces in vlp23, colour-coordinated with **d**. The black arrows point to the frustration states of the interfaces in **a**, **b**, and **c**, respectively. **f**, The frustration state and the number of residue pairs involved in the helix interactions are correlated – the more interactions, the lower the frustration state.

The distribution of local frustration across all dimer interfaces in vlp23 is summarised in Fig. 4e, where the black arrows (a-c) point to the frustration states of the interfaces in Fig. 4a-c. Interfaces with the top 50 highest frustration values are highlighted in red, and the next 51-100 in purple. The corresponding interfaces are mapped onto the capsid structure as coloured spheres (Fig. 4d).

Strikingly, in addition to the concentration of frustration in the two end caps, which are dominated by seam lines, the most highly frustrated interfaces in the capsid’s midsection co-localise with the seam side of the data-derived lattice identified by the geometric index (Fig. 4d). In contrast, mapping the 50 lowest-frustrated interfaces reveals no distinct spatial clustering (Supplementary Fig. 7a-b). The colocation of frustration with the seam line of the CA lattice suggests that biophysical properties of the HIV-1 cone are strongly coupled to the underlying geometric organisation of its CA lattice.

The correlation between the local frustration state and the proportion of water-mediated contacts further suggests that the geometric organisation may modulate function by influencing interfacial solvation, which contributes to frustration differences. To test the robustness of the observed seam-frustration relationship, we repeated the analyses excluding all water-mediated contacts. The highly frustrated interfaces remained localised at the seam side (Supplementary Fig. 8a), and the correlation between frustration and contact number was preserved (Supplementary Fig. 8b-e), showing that the observed relationships do not merely depend on solvent-mediated interactions.

Together, these results highlight the importance of selecting a geometric model that not only captures the correct surface topology but also reproduces the intrinsic biophysical heterogeneity of the HIV-1 capsid.

### The geometric index distinguishes between HIV-1 capsid reconstruction protocols

The seam line-associated frustration discussed above may also play a role in capsid assembly or disassembly. Here we use the geometric index to analyse if the reconstruction protocol applied to the cryo-EM data leaves imprints on the atomistic model. We perform a comparative analysis of three capsid models (vlp23 Fig. 5a, PDB: 3J3Y Fig. 5b, and PDB: 3J3Q Fig. 5c): a fullerene, a Kagome, and a data-derived surface lattice model. The distributions of their geometric indices are shown as bar charts (Fig. 5), which act as fingerprints of the models. The density lines (purple for the fullerene, green for the Kagome, and red for the data-derived surface lattice) are plotted together with histograms that distinguish the geometric indices surrounding hexamers (orange) and pentamers (cyan). Corresponding pictorial representations of the characteristic triangles (grey surrounding hexamers, cyan surrounding pentamers) are displayed on the right in frames coloured to match the respective density lines.

**Fig. 5.**
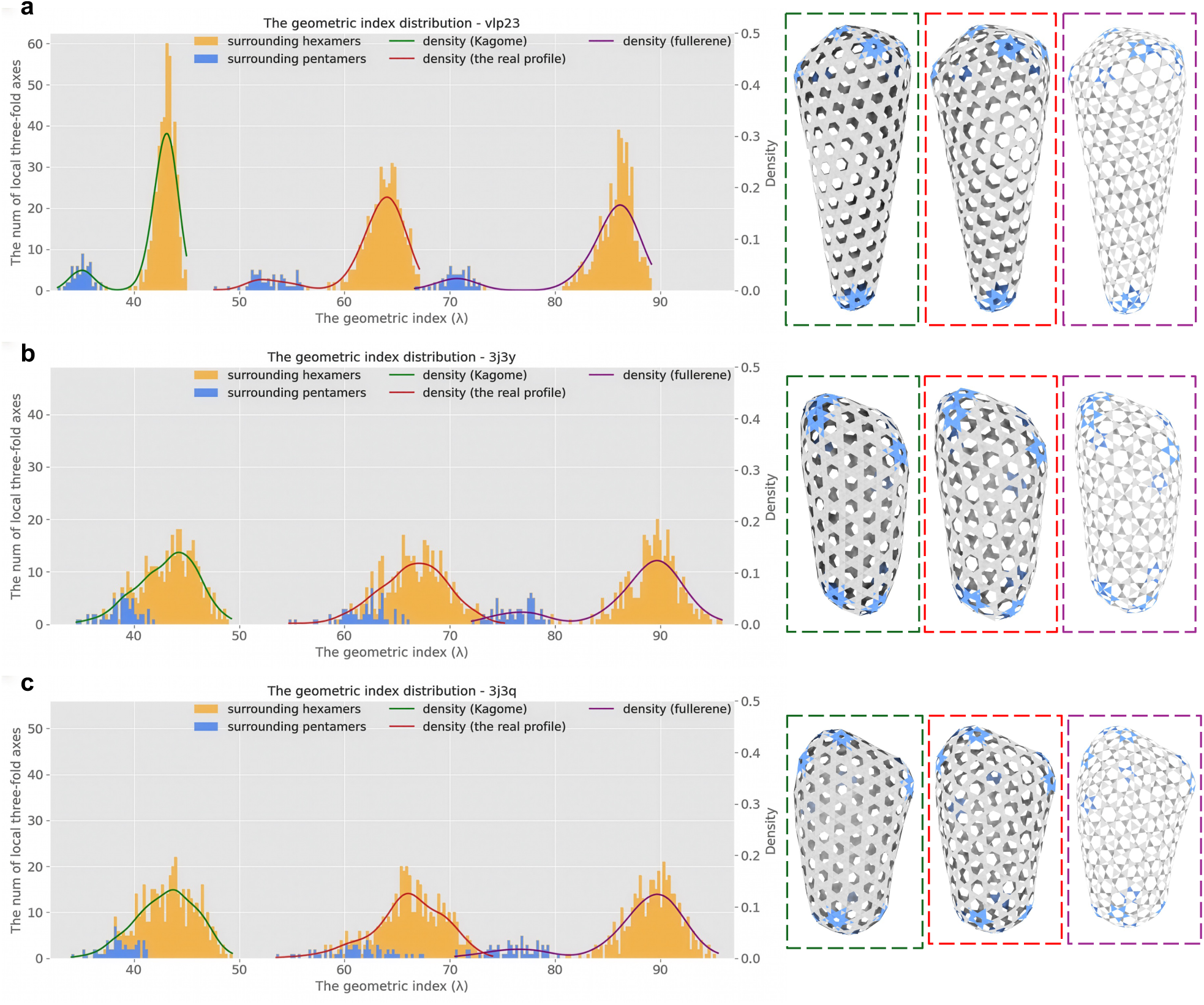
Data-derived models have distinct geometric index distribution profiles for different assembly scenarios. Left: the distribution of the geometric indices for the different geometric representations of (**a**) vlp23 and (**b**) PDB: 3J3Y (**c**) PDB: 3J3Q. Here the real profile represents our data-derived model. The geometric index distribution of the data-derived model is bi-modal in vlp23 but merged into a single peak in PDB: 3J3Y and 3J3Q. Right: capsid representation in terms of characteristic triangles (grey surrounding hexamers, cyan surrounding pentamers) for the fullerene lattice architecture (green dashed frame), Kagome lattice (purple dashed frame), and the data-derived model (red dashed frame).

For all capsids, the fullerene lattice exhibits a bimodal geometric index distribution, consistent with the behaviour observed in our computational carbon-fullerene cones. However, striking differences emerge in the data-derived and Kagome lattices. In vlp23 (Fig. 5a), both lattices retain bimodal distributions, whereas in 3J3Y (Fig. 5b) and 3J3Q (Fig. 5c) the two peaks are merged into a single negatively skewed mode.

These differences primarily reflect the distinct reconstruction and simulation protocols used to build each model. In vlp23, the complete capsid was first assembled by fitting capsomers (hexamers/pentamers) into a cryo-EM density map along a Hamiltonian path, followed by molecular dynamic (MD) relaxation of the entire structure^18^. In contrast, 3J3Y and 3J3Q were constructed through MD simulations applied initially to isolated patches composed of hexamers and pentamers, which were later concatenated onto a fullerene backbone and further refined by MD^16^. The early relaxation of local patches likely allowed the geometry to adapt toward a local equilibrium before global assembly. Although the present analysis is based on static reconstructed structures rather than explicit dynamic assembly trajectories, it shows that the geometric index is a quantitative descriptor of the final lattice state that may reflect how capsomers equilibrate under different geometric constraints during capsid formation or reconstruction. In this sense, the observed differences are consistent with the hypothesis that assembly kinetics and relaxation process influence capsid geometry.

In line with this interpretation, Supplementary Fig. 9 shows that hexamers in vlp23 remain close to regular hexagonal geometry, whereas those in 3J3Y and 3J3Q display marked distortions, as quantified by differences in their main diagonals. These contrasting reconstruction pipelines – global reconstruction followed by relaxation versus pre-equilibrated patch reconstruction – produce distinct geometric organisations, and potentially different biophysical properties of the resulting capsids. Although these structural variations originate from the modelling routines rather than direct physical processes, they provide insights into how alternative assembly scenarios might influence capsid architecture. Finally, mapping the geometric index orientations *v*^↔^ onto the data-derived models (Supplementary Fig. 10) reveals some specific patterns, such as circular orientations consistent with the pre-assembled patches in 3J3Y and 3J3Q used in the reconstruction. This further supports the hypothesis that capsid assembly pathways and protocols leave measurable geometric imprints on the final topography that can be captured using the geometric index.

### The geometric index captures local curvature and lattice distortions

To further explore the geometric information captured by the geometric index, we examined its relationship with local curvature in the data-derived model of the HIV-1 capsid vlp23. Fig. 6a illustrates seven neighbouring hexamers, each coloured distinctly, with four local threefold axes marked by characteristic triangles (grey). The geometric index of the central triangle is highlighted in magenta. We defined two curvature factors – the angles φ_1_ and φ_2_ – to quantify local surface bending (Fig. 6b). The two purple vectors (see top of Fig. 6b) represent the surface normal vectors of the triangles adjacent to the geometric index of the central triangle. The first curvature factor, φ_1_, is defined as the angle between these two normal vectors. The black vector (bottom of Fig. 6b) denotes the normal of the central triangle, while orange vectors correspond to the normal vectors of its three surrounding triangles. The second curvature factor, φ_2_, is defined as the mean angle between the orange and black vectors. The curvature measurements used here provides a coarse-grained measure of surface bending that integrates contributions from multiple capsomer interfaces. Unlike dihedral angles used in icosahedral assembly studies^24,45^, which describe pairwise rotations between adjacent subunits in a fully triangulated lattice, the measurements used here probes curvature within the local environment of adjacent hexamers in the native gyrated hexamer-pentamer topology without redefining the lattice into triangular elements.

**Fig. 6.**
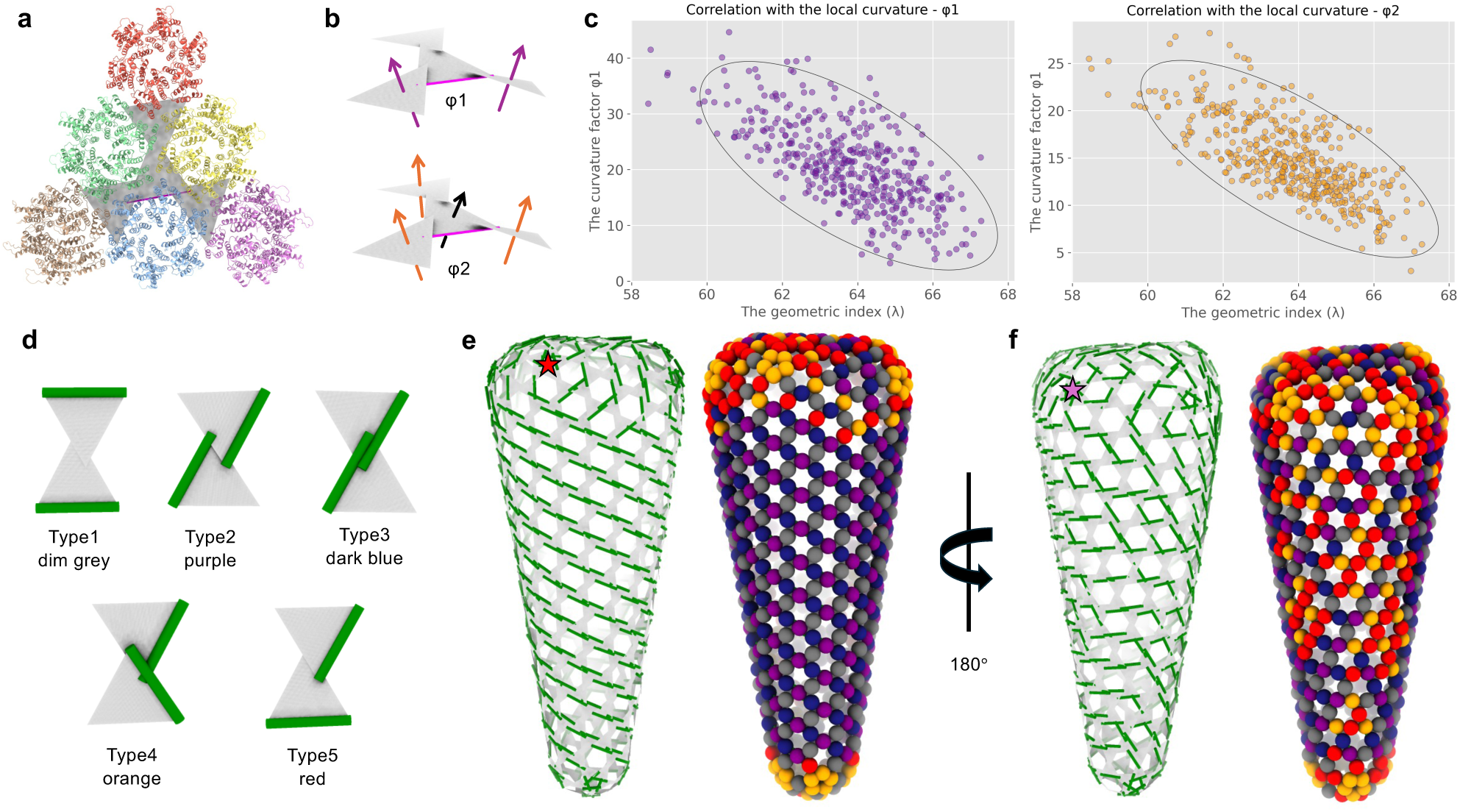
The geometric index of the data-derived model correlates with local geometric features. **a**, Four characteristic triangles (grey) corresponding to the six CA hexamers shown in different colours; the geometric index of the central triangle is indicated in magenta. **b**, The four characteristic triangles in **a**. Top: the two purple arrows represent the normal vectors of the two triangles connected by the line representing the geometric index of the central triangle. The angle between the two purple vectors is the curvature factor φ1 in **c**. Bottom: the three orange arrows represent the normal vectors of the three triangles surrounding the central triangle, and the black arrow is the normal vector of the central triangle. The average over the angles between the black vector and the three orange vectors, respectively, is the curvature factor φ2 in **c**. **c**, The geometric index correlates with the curvature factors φ1 (left) and φ2 (right) and is thus a suitable indicator of capsid geometry. **d**, Five types of local geometric organisations defined by the orientations of neighbouring geometric indices (green lines); types 1-3 are parallel, whereas types 4-5 are non-parallel. **e**-**f**, Characteristic triangles (grey) and geometric indices (green) on opposite sides of the capsid (red and light purple stars). Each pair of neighbouring threefold axes is colour-coded – dim grey (type 1), purple (type 2), dark blue (type 3), orange (type 4), and red (type 5). Parallel types from sheet-like regions on the red-star side, while non-parallel types cluster on the light-purple-star side and at both termini, coinciding with highly frustrated interfaces (Fig. 4).

As shown in Fig. 6c, the length of the geometric index (λ) is inversely correlated with local curvature: smaller geometric indices correspond to regions of higher curvature. This finding confirms that the geometric index not only encodes lattice organisation but also serves as a quantitative indicator of local geometric deformation across the capsid surface.

In addition to its length, the orientational patterns of the geometric indices *v*^↔^ are important to capture more detailed topographic features (Fig. 6d). Neighbouring geometric indices, represented by green lines, were grouped into five distinct types based on their relative orientations. In types 1-3, the indices are parallel, while in types 4 and 5 they are non-parallel. Fig. 6e and 6f (left) show pictorial representations of characteristic triangles (grey) and their geometric indices (green) on opposite sides of the capsid, denoted by the red and light purple stars, respectively. Each pair of neighbouring threefold axes was assigned one of the five types and colour-coded on the capsid surface – dim grey (type1), purple (type2), dark blue (type 3), orange (type 4), and red (type 5). The right panels of Fig. 6e and 6f (see also Movie 4) depict the corresponding twofold axes as coloured spheres.

The parallel configurations (type 1-3) predominate on the side marked by the red star, forming sheet-like surface regions, whereas non-parallel configurations (types 4-5) are concentrated on the opposite side (with the light purple star) and at both termini of the capsid. The accumulation of type 4 and 5 configurations at the highly frustrated dimer interfaces identified in Fig. 4 indicates a coupling between local topography and molecular frustration.

Together, these analyses establish that the geometric index – through both its length and orientation – provides a quantitative framework for describing capsid curvature and surface organisation. In the following section, we apply this framework to characterise the geometric features of cofactor and small-molecule binding regions and use this to predict putative CypA binding sites.

### The geometric index reveals structural signatures at cofactor binding sites in tubular assemblies

Host cofactors interact with the HIV-1 capsid throughout the viral life cycle, coordinating processes such as trafficking, nuclear import, and uncoating^3^. Two primary binding interfaces have been identified on the capsid surface^35–37^. The first, a flexible loop between helices 4 and 5 in the CA N-terminal domain (CA_NTD_), binds cytoplasmic cyclophilin A (CypA)^30,31^ and the cyclophilin domain of nuclear pore component Nup358^46,47^. The second, a pocket at the interface between CA monomers, engages FG motifs from host proteins such as Nup153, Sec24C, and CPSF6^48–50^, and accommodates small therapeutic molecules like lenacapavir (LEN)^32–34^. Not all such interfaces are occupied *in situ*, and the geometric determinants of these binding events remain poorly quantified. To address this, we applied the geometric index as a local descriptor to characterise capsid surface organisation at the cofactor and drug molecule binding regions.

We analysed experimentally available data sets (PDB: 6Y9W, 6Y9V, 6Y9X, 6Y9Y, 6Y9Z, and 6ZDJ) for six CypA-CA complexes of tubular assemblies exhibiting different helical symmetries. Monomers directly bound to CypA via the canonical CypA-binding loop were identified and coloured in cyan (Figs. 7a&b and Supplementary Fig. 11). Two monomers located around the same three-fold axis are coloured in lime and yellow, respectively. Characteristic triangles were constructed based on the N195 Cα in the C-terminal domains (CA_CTDs_). The geometric index of each triangle (dashed cyan line) describes the local relationship between adjacent capsomers. Across all six complexes, the CypA-binding monomer consistently lies opposite the geometric index (the shortest edge) of the characteristic triangle, and the geometric index of the CypA-associated triangle is parallel to, but smaller than, that of its neighbouring triangle (purple dashed lines) – a configuration corresponding to the type 2 dimer interfaces defined in Fig. 6d. This consistent pattern suggests that preferred CypA-binding sites coincide with type 2 configurations in the tubular lattice.

**Fig. 7.**
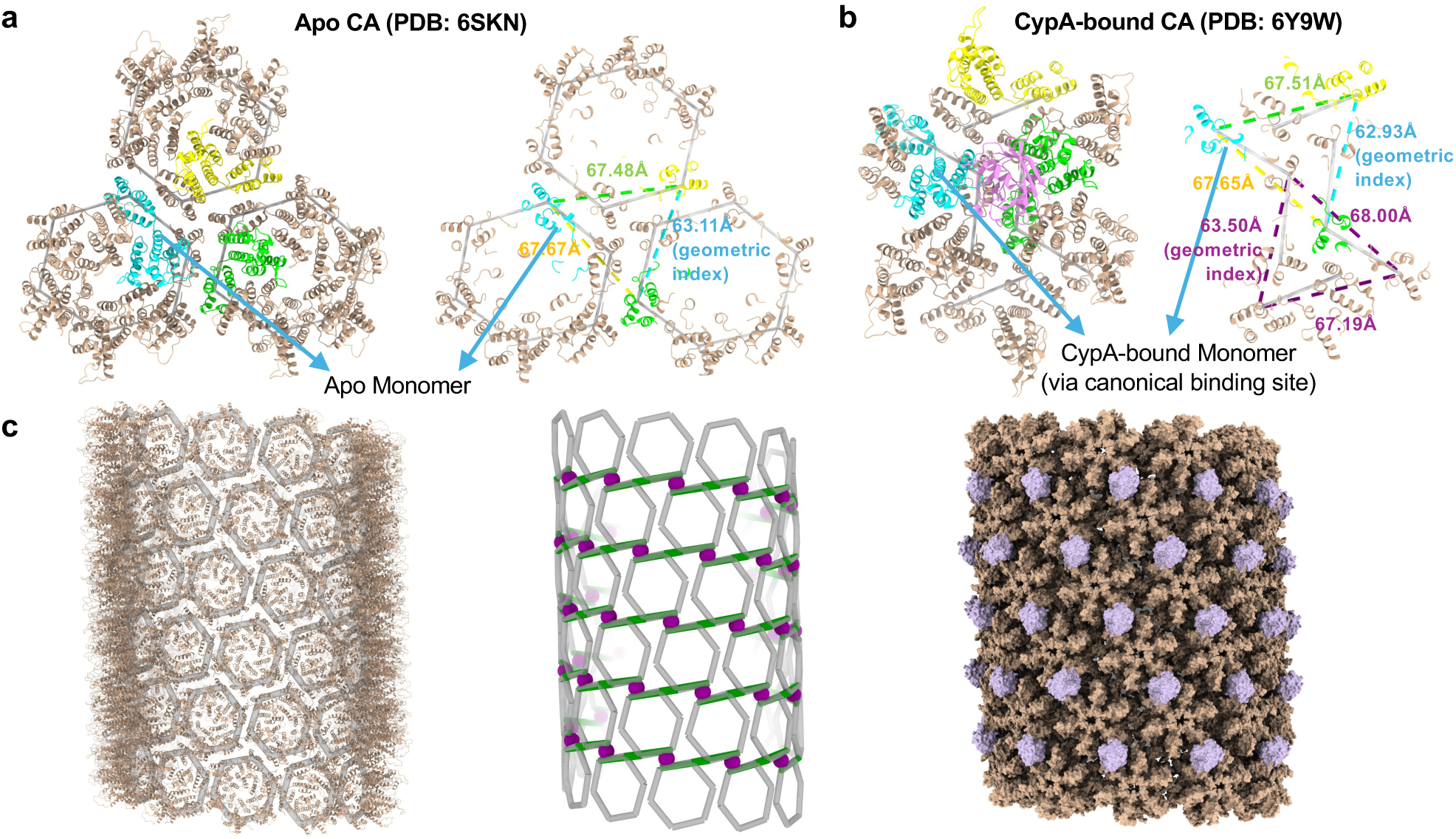
The geometric index as an indicator of cofactor (CypA) binding. **a**, The Apo CA complex in a tubular structure with helical symmetry (−13, 8) (PDB: 6SKN), where the monomer coloured in cyan is matched to the corresponding CypA-binding monomer (cyan) in **b** by the Needleman-Wunsch algorithm. Right: the three dashed lines indicate the characteristic triangle, with numbers denoting their lengths in the corresponding colours. **b**, A CypA-CA complex in a tubular structure with helical symmetry (−13, 8) (PBD: 6Y9W), where CypA is coloured in magenta. Right: six dashed lines construct two characteristic triangles at two neighbouring local three-fold axes. **c**, CypA-binding site prediction based on the features revealed in **a** and **b**. Left: an HIV-1 capsid tube (PDB: 6X63) in brown with the data-derived profile of CA hexamers in light grey. Middle: the data-derived geometric model, with green lines representing the geometric indices. Purple spheres predict CypA-binding sites. Right: CypAs in light purple are mapped onto the predicted binding sites of the capsid tube (PDB: 6X63), by aligning the CypA-binding monomer in the complex (PDB: 6ZDJ) in Supplementary Fig. 11e to the corresponding CypA-binding monomers in the capsid tube.

Mapping the geometric indices onto the surface of a CA tubular assembly^51^ (PDB: 6X63, helical symmetry −12, 11) further supports this interpretation (Fig. 7c, left and middle). Only parallel configurations (type 1-3) are present, with no distorted (type 4-5) arrangements, indicating a continuous sheet-like topology without seam lines. When CypA-binding sites are overlaid as purple spheres at the type 2 interfaces, the smaller geometric index identifies the canonical CypA-binding monomer. Alignment of the CypA-CA complex (PDB: 6ZDJ, symmetry −13, 10) onto these positions yields a model (Fig. 7c, right, and Movie 5) in which bound CypAs follow one of the three characteristic orientations in the hexameric surface lattice.

Earlier work^25^ suggested that CypA selectively binds the CA hexamer array direction with the highest curvature. However, their more recent high-resolution data^27^ combining cryo-EM density and atomic models indicate that CypA instead bridges two CA hexamers along the direction of lowest curvature, with additional CypA molecules occasionally bound adjacent to these at the trimer interfaces. The Apo and CypA-CA complexes analysed here (Figs. 7a,b; Supplementary Fig. 11) derive from these updated data^27^, and our predicted CypA-binding sites in the CA tubular assembly^51^ (PDB: 6X63) are consistent with the hexamer array orientation with the lowest curvature.

In tubular assemblies, the three hexamer array orientations can be described as most curved, least curved, and intermediate, depending on the helical symmetry. However, absolute curvature varies between different helical symmetries, making it difficult to determine whether these three relative directions are directly comparable across different tubes – particularly considering capsomer tilt, twist^26^, and internal curvature^27^. We therefore further quantified the curvature patterns along the three hexamer array orientations (ori1-ori3) in tubular assemblies with helical symmetry (−13, 8; PDB: 6SKN) and (−12, 11; PDB: 6X63) using a covariance matrix analysis (see Methods). As illustrated in Supplementary Figs. 12 and quantified in Supplementary Fig. 13, curvature strengths differ between symmetries. However, their relative geometric patterns remain conserved, with the three relative curvature directions consistently aligning with the geometric index configurations of types 1-3 (Fig. 6d), respectively. These results suggest that the geometric index captures relative local curvature patterns across different protein assemblies, as revealed by the covariance matrix approach, thereby refining our understanding beyond merely absolute curvature values alone.

### Ligand-specific curvature and frustration responses revealed by the geometric index

To examine how ligand binding influences local geometry, we first compared Apo and CypA-bound CA complexes with identical helical symmetry (Apo CA, Fig. 7a, PDB: 6SKN; CypA-CA, Fig. 7b, PDB: 6Y9W). Alignment of the corresponding monomers revealed a decrease in geometric index upon CypA binding. According to the anti-correlation between the geometric index and local curvature derived in the previous section (Fig. 6c), this should be indicative of a local increase in curvature. Prior experimental studies show a correlation between increasing CypA concentration and progressively higher curvature in the assemblies, with examples ranging from long tubes to cones and spherical particles^25^, consistent with the enhanced curvature upon CypA binding. To further validate this geometric interpretation, we quantified local curvature by measuring the angle between the normal vectors of two adjacent triangles (purple triangles in Supplementary 15a&b) before and after cofactor binding. The angles increased after CypA binding, confirming the hypothesised local curvature increase with decreasing geometric index.

Equivalent analyses for LEN- and CPSF6-peptide-bound CA complexes (Supplementary Fig. 14), derived from CA lattice structures, showed the opposite trend – an increase in geometric index, that we hypothesize to be an indicator of local flattening of the surface lattice. We again validate this via curvature measurements based on two adjacent triangles. Curvature decreased after ligand binding (purple triangles in Supplementary 15c-f), consistent with our interpretation of the geometric index. Although studies on CPSF6-induced geometric changes are limited, recent reports^34,52^ indicate that LEN binding causes lattice flattening, defective capsid closure, and loss of essential pliability, in agreement with our results.

We next investigated whether ligand-induced geometric changes are coupled to molecular frustration at both the binding sites and at the adjacent dimer interfaces (see Methods). Comparison of structural data before and after CypA binding (Fig. 8a-d, see also Movie 6 & 7) shows that the canonical CypA-binding site is located above the dimer interface, whereas CPSF6 binds adjacent to the interface itself (Fig. 8e-h). As expected, both ligands reduce frustration at their respective binding regions. Notably, however, the dimer interfaces exhibit opposite responses: CypA binding increases frustration at nearby interfaces, as indicated by a lower average frustration index, a higher proportion of highly frustrated contacts and a reduced proportion of minimally frustrated contacts (Tab. 1 & 2), whereas CPSF6 binding decreases interfacial frustration, reflected by a higher average frustration index and a reduced proportion of highly frustrated contacts (see Methods). These contrasting responses are consistent with the observed geometric effects – curvature enhancement associated with increased interfacial frustration for CypA, versus lattice flattening associated with reduced frustration for CPSF6.

**Fig. 8.**
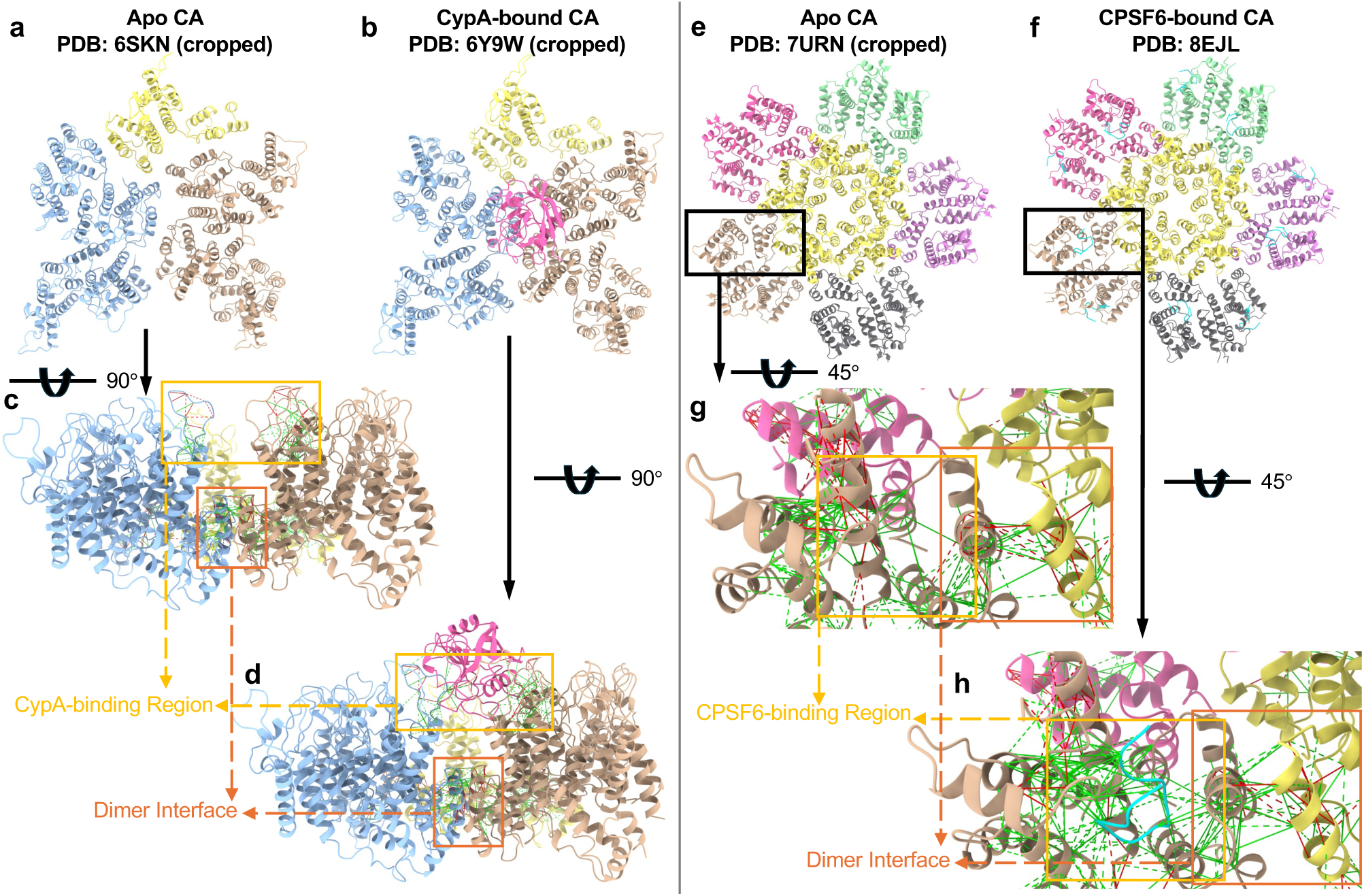
Molecular frustration at binding sites and dimer interfaces before and after cofactor binding. All complexes were cropped from original PDB structures to ensure consistent molecular frustration comparisons. **a**, Apo CA complex prior to CypA binding, with three regions from different hexamers highlighted in distinct colours. **b**, CypA-CA complex, with CypA shown in hot pink. **c**-**d**, Views rotated by 90° relative to **a** and **b**, respectively, highlighting the CypA-binding region (black frame) and dimer interface (dark orange frame). Highly (red) and minimally (green) frustrated molecular interactions are shown, while neutral interactions are omitted for visibility due to their abundance. **e**, Apo CA complex prior to CPSF6 binding, with six regions from different hexamers and pentamers highlighted in distinct colours. **f**, CPSF6-CA complex, with CPSF6 shown in cyan. **g**-**h**, Enlarged views rotated by about 45° of the regions indicated by black frames in **e** and **f**, respectively, showing the CPSF6-binding region (black frame) and dimer interface (dark orange frame), with molecular interactions superimposed.

**Tab. 1.** Changes in average configurational frustration index upon cofactor binding. The reported index values correspond to the configurational frustration index (Z-score). Lower values indicate higher frustration, whereas higher values indicate lower frustration.

|  | CypA binding |  |  | CPSF6 binding |  |  |
| --- | --- | --- | --- | --- | --- | --- |
|  | Before | After | Num. of res. | Before | After | Num. of res. |
| **Binding region** | -0.050 | 0.296 | 34 | 0.089 | 0.387 | 425 |
| **Dimer interface** | -0.019 | -0.127 | 84 | -0.005 | 0.134 | 140 |

Residue-wise frustration analyses across the binding monomers further support this coupling (see Methods and Supplementary Figs. 16 and 17). CPSF6 binding reduces frustration at both the binding region and at the adjacent dimer interfaces, whereas CypA binding lowers frustration at the binding site but increases frustration at the nearby interface. These results suggest that cofactor binding can have an allosteric effect on nearby interfaces that are not in direct contact with the binding site.

While these observations highlight opposing geometric and frustration responses at the two primary binding sites – curvature enhancement upon CypA binding versus surface flattening upon FG-motif or LEN binding – they should be interpreted cautiously. The CypA complexes originate from tubular assemblies, whereas the LEN and CPSF6 complexes are derived from CA lattices; thus, part of the observed differences may arise from the structural context rather than ligand binding alone. Moreover, cofactor binding is likely to affect not only local geometry but also biochemical and mechanical properties of the lattice.

Nevertheless, the relationship between our geometric index, curvature at the dimer interfaces and molecular frustration, in line with experimental observations, supports a genuine site-specific coupling between ligand binding, local frustration, and geometric modulation. These findings suggest that cofactors and inhibitors may regulate capsid function by exploiting local geometric flexibility along distinct structural axes. These links between geometry and biophysical function can be exploited in the design of virus-like particles for a host of applications in biotechnology and medicine. Implementing these geometric constraints in the context of machine learning techniques^53^ can be used to achieve desired properties in lentiviral vectors. For example, designing appropriate seam lines in these capsid architectures can afford predictive control over cargo release mechanisms.

## Discussion

The geometric index introduced here acts like a mathematical microscope, providing a quantitative link between the local surface lattice architecture of an HIV-1 capsid and its biophysical properties. By capturing both the size and orientation of local lattice distortions, it connects atomic-scale geometry with macroscopic features such as curvature, molecular frustration, and cofactor binding, revealing a deep coupling of capsid geometry and function. A surprising conclusion from our study is that the choice of surface lattice in an HIV capsid model is paramount for capturing essential aspects of its biology. In contrast to the conventionally used fullerene lattice, only the data-derived surface lattice links potential seam-line locations with local frustration patterns. In addition to this, the geometric index can be used to identify curvature-dependent binding of cofactors and small molecules, and to predict the geometric and biophysical responses to these binding events. Overall, the geometric index provides a unifying descriptor of HIV-1 capsid geometry that connects its local geometric organisation with variations in curvature and molecular frustration. While the geometric index provides a minimal descriptor of lattice distortion, future extensions could incorporate strain-like measures or elastic parameters to provide a more direct mechanical interpretation of lattice deformation along the lines of previous studies in other viral families^54^. Extending this framework to other retroviruses and to artificial capsid systems may uncover universal geometric rules of viral architecture.

Beyond these structural insights revealed by our study, the geometric index opens up new avenues for the study of capsid assembly and evolution. The distinct geometric signatures observed across capsids reconstructed via different reconstruction protocols suggest that the assembly methods used may leave characteristic imprints on capsid structure. Exploring the implications of this for the assembly kinetics of different cone structures through the lens of the geometric index may clarify how HIV-1 controls its final morphology, particularly in light of the experimentally reported variations in pentamer distributions^55^. The role of the viral genome also warrants attention in this context. While capsid proteins can assemble into cones or tubes *in vitro* in the absence of RNA^56^, *in vivo* data show that cylindrical shells rarely encapsulate genomes, whereas conical ones consistently do^17,57^. This observation implies that genome-capsid coupling could influence cone geometry, potentially guiding the formation of functionally competent cores. Our geometric index can be used to quantify any curvature effects that may occur in the presence and absence of the genomic RNA, providing a platform for studying how interactions with viral genomes may bias assembly outcomes.

The geometric principles uncovered here may also shed new light also on antiviral and synthetic design strategies. In analogy to Capsid Assembly Modulators (CAMs) developed for Hepatitis B virus^58,59^, geometric features derived from our criterion could guide the development of HIV-1 assembly inhibitors that destabilise or misdirect core formation at pivotal locations in the capsid surface revealed by the geometric index. For example, modulating the sizes or orientations of characteristic triangles via small molecular weight compounds might bias assembly towards non-productive or malformed structures. In addition, the theory presented here can be harnessed in gene therapy, where HIV-1 cores serve as lentiviral vectors for payload delivery. By implementing the structural constraints embodied by the geometric index in the context of machine learning techniques, capsid proteins can be engineered to improve vector stability or cargo capacity. Such geometry-informed, AI-driven molecular design approaches offer promising avenues for antiviral viral vector engineering.

## Methods

### Computational Fullerene and Quantum Mechanical Methodology

#### Fullerene

Following a previous publication^16^, the Fullerene^38–41^ program was used to create multiple fullerenic models of the HIV-1 capsid. Specifically, the ring spiral pentagon indices (RSPIs)^60,61^ corresponding to the broader end (the last seven indices) were systematically altered by increments of ±1, ±2, and higher values. For this purpose, input files for the geometry search were generated outside of the program Fullerene using a template input text file modified with sed. Each modified RSPI was then tested within Fullerene to determine whether it produced a valid closed fullerene. This validation was performed without attempting to generate a three-dimensional geometry, relying instead on the program’s ability to verify consistency of the candidate RSPI with the specified number of carbons and the canonical RSPI rules. The majority of RSPI variations proved invalid. Therefore, searches were conducted in parallel to efficiently identify the small subset of valid candidates. A valid RSPI indicates the existence of a closed three-dimensional geometry consistent with the indices.

Candidate RSPIs that passed validation were used as input for three-dimensional embedding procedures available in Fullerene, including the 3D Tutte^62^ embedding method. Multiple embedding algorithms were tested because some methods yielded less deformed geometries than others; we found that 3D Tutte was more consistent at producing closed geometries than the adjacency matrix eigenvector (AME) method^63^. To improve reproducibility and reduce bias, the embedding process was automated and repeated with different methods. For each candidate RSPI, embedding parameters were optimised to obtain satisfactory initial three-dimensional structures. Although log files provided preliminary indicators of structural quality, visual inspection was used to confirm the adequacy of the resulting geometries during parameter optimisation.

Subsequent structural optimisation was carried out using semi-empirical quantum mechanical methods implemented in MOPAC^64^ with Parametric Method 3 (PM3)^65^, PM6^65^, and PM7^65^ Hamiltonians. Final refinement of carbon fullerene geometries was performed with higher-level quantum mechanical approaches based on density functional theory (DFT) as described below. As a result, there are three different topological classes of vertices 352, 452, and 524. These structures correspond to fullerenes with 352 (C_352_), 452 (C_452_), and 524 (C_524_) carbon atoms, which contain 166, 216, and 240 hexagonal faces, respectively, in addition to the 12 pentagonal faces required by topology. Each fullerene has multiple possible isomers that arise from different arrangements of the pentagonal and hexagonal rings on the carbon cage. The full data set contains 654 fullerene structures divided into 325 of C_352_, 39 of C_452_, and 290 of C_524_.

#### Optimisation

To investigate the energy landscape of the preferential shape of fullerene-based capsid topologies we treated the vertices of the generated fullerene cones as carbon atoms and subjected them to tight-binding (TB) density functional theory (DFT) optimizations and final energy evaluations. This allowed us to estimate the optimal fullerenic shape-per-discrete vertices count. In order to optimise the fullerene structures, we employed the Orca 6.0^66^ quantum chemistry package. To determine the minimum energetics of each fullerene cone we used the GFN2-xTB DFT tight binding approach of the Grimme group^64^. While more accurate energies would likely be obtained through more sophisticated wavefunction approaches, or even through canonical DFT, given the size of the dataset and atomic count of each fullerene cone, the GFN2-xTB approach provides an efficient and sufficiently accurate metric for ranking^64^.

The binding energy per atom derived from the quantum mechanical calculations is defined through the following equation

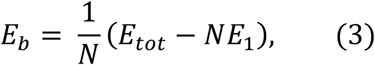

where *E*_tot_ is the total energy of the fullerenic cone from GFN2-xTB calculations, *E*_1_ is the energy of a single carbon atom, *E*_b_ is the binding energy, and *N* is the number of carbon atoms in the fullerene.

#### Capsid Structures

To determine topologically favourable capsid structures based on the analysed fullerene cones we have introduced two criteria for characterisation of cone structures. The first criterion is related to the binding energy per atom, as described by equation 3, and the second characterizes the spherical asymmetry of the structures, as presented by Diaz-Tendero et al^67^.

The binding energy of a fullerene cone is closely related to the geometrical distortion of its surface. This has been revealed by Solov’yov et al^68^ by demonstrating that the binding energy of a single wall carbon nanotube of arbitrary chirality can be calculated with a 99 % accuracy as a sum of surface, curvature and edge energy terms. Because of the similarity of their model and those used to study liquid droplets, nuclei and clusters, they adopted the name liquid surface model (LSM)^68^.

One can generalize the LSM to the energetics of the fullerene cones studied here. In the fullerene case the dominant contributions to energy arise due to the surface area of the cone and the curvature of its surface. According to the LSM the lower values of the binding energy defined in equation 3 suggest the corresponding fullerene structures have fewer surface distortions, i.e., the overall shape is smoother, and the surface is less curved. Following this argument, in the present analysis we have chosen the structures with the lower *E*_%_ value as those being more compact, and, therefore, topologically favourable.

Binding energy provides a natural means to characterize the quality of the fullerene cones. We have introduced a second parameter ξ which describes the asymmetrical components and is used in conjecture with the binding energy *E*_%_ to highlight favourable structures. We can define the cone asymmetry through the following equation, adapted from Díaz-Tendero et al^67^:

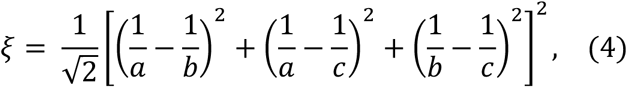

where, *a*, *b*, and *c* are the normalised principal components *I*_xx_, *I*_yy_, and *I*_zz_ of the moment of inertia tensor of the cone, respectively, as defined by equations 5-7.

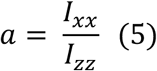

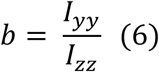

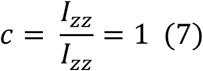

We orientate the axes, so that the z-component of the inertial tensor is the largest. Given the oblong cone shape of the resultant fullerenes, *a* < *c* and *b* < *c* hold for all configurations. The asymmetry parameter ξ in equation 4 describes the deformation deviation from the fullerene cone compared to a sphere of radius *R* = *a* = *b* = *c* (corresponding to ξ = 0). For an axially symmetric ellipsoidal shape *a* = *b* = *R* < *c*, we have

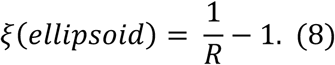

From this equation, it follows that for *R* = ½, ξ = 1. The correlation between the asymmetry parameter ξ and the energy *E*_b_ is illustrated in Supplementary Fig. 2.

### The Construction and Comparison of Geometric Representations

To construct the data-derived geometric model, we use the N195 Cα in the CA_CTD_ of each monomer as a reference point. This site was chosen because the distance between diametrically opposed N195 Cα (orange dashed lines in Fig. 3d) correlates with the internal curvature of the CA hexamers^27^. As illustrated in Fig. 3d, profiles of three CA hexamers (grey hexagons) derived from the N195 Cα positions show that their hexagonal edges are not perfectly coincident, leaving distinct inter-hexamer gaps.

For comparative analysis, we generated fullerene and Kagome lattices (Supplementary Fig. 3). In these two lattices, the fullerene vertices (black dots) were defined as the centroids of triplets of the neighbouring hexamer vertices (light purple arrows), while the Kagome lattice (grey dots) were defined as midpoints between corresponding vertices of adjacent hexamers located across the local twofold axes (brown arrows). Each Kagome vertex thus represents a pair of monomers connected through a dimer interface.

The three geometric representations – the data-derived, fullerene, and Kagome lattices – are compared in Supplementary Fig. 4. All models were constructed onto seven uniformly sized hexagons with different colours. The fullerene lattice produces slightly larger hexagons covering the small gaps between the coloured hexagons (Supplementary Fig. 4b), while the Kagome lattice reproduces the same hexagon size but rotated by approximately 30° around the hexagons’ centres (Supplementary Fig. 4d&e). Comparisons of the data-derived lattice (represented by the coloured hexagons) with the fullerene and Kagome lattices are shown in Supplementary Figs. 4c&f, respectively.

To evaluate geometric discrepancies, we superimposed the three lattice types on representative regions of the HIV-1 capsid model vlp23. Supplementary Fig. 5 shows the corresponding overlays of the data-derived model (grey lines) with the fullerene (black dots), and Kagome (grey dots) lattices. The characteristic triangles that define different geometric indices were extracted from these overlays, allowing a direct comparison of the data-derived lattice (orange dashed lines) with the fullerene and Kagome lattices (purple dashed lines) in Supplementary Fig. 6.

### Molecular Frustration Analysis

Local frustration was quantified using the configurational frustration index of protein contacts computed with the Protein Frustratometer^28,29^, which evaluates the energetic contribution of native residue-residue contacts relative to an ensemble of alternative (decoy) interactions. The configurational frustration index was chosen because it correlates strongly with protein-protein binding interfaces^42,43^. For each pair of residues *i* and *j*, a configurational frustration index (Z-score) was computed to assess how favourable the native interaction energy is compared to randomised contacts, including electrostatic contributions. Based on that, contacts were classified as highly frustrated (frustration index ≤ −1), neutral (−1 < frustration index < 0.78), or minimally frustrated (frustration index ≥ 0.78). These thresholds follow established conventions in previous studies^28,29^.

### Frustration at Dimer Interfaces across the vlp23 capsid surface

To evaluate interfacial frustration, we defined a local frustration parameter (γ_i_) at each dimer interface *i* as the average frustration index across all contacts involving the H9 helices:

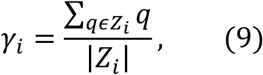

where *Z*_i_ represents the set of frustration indices for all contacts associated with the H9 helices at the *i*^th^ dimer interface. To ensure consistency with the dimer baseline structure (PDB: 2KOD), residues 146-221 from each monomer (76 residues per monomer) at every dimer interface were included in the frustration computation. These residues encompass the H9 helices and neighbouring regions that mediate inter-monomer contacts.

### Frustration Changes upon Cofactor Binding

For ligand-binding analysis, residues interacting with cofactors (CypA and CPSF6) were identified from the corresponding structural complexes (PDB: 6Y9W and 8EJL). In the CypA-bound structure, 34 residues participate in interactions with CypA, as shown in Tab. 1 (see Source Data file for details). By contrast, the CPSF6-bound structure contains five equivalent binding regions surrounding the central pentamer, each comprising 85 residues (425 residues in total; Tab. 1; see Source Data file for details).

To assess coupling with nearby interfaces, we additionally analysed adjacent dimer interfaces. Each interface involves two H9 helices (residues 179-192) from the contacting monomers (28 residues per interface). In the CypA-bound complex, three adjacent interfaces are available for comparison (84 residues total; Tab. 1), whereas in the CPSF6-bound complex, five interfaces surrounding the pentamer are analysed (140 residues total; Tab. 1).

The local frustration at the binding regions and interfaces was then computed using Equation (9), where *Z*_!_ denotes the set of frustration indices for all contacts within a 5 Å neighbourhood of the selected residues (Tab. 2).

**Tab. 2.** Changes in the proportion of highly and minimally frustrated contacts upon cofactor binding. Proportions are calculated for contacts (bonds) within a 5 Å neighbourhood of residues in the selected regions before and after ligand binding.

|  | CypA binding |  |  |  |
| --- | --- | --- | --- | --- |
|  | Before | Num. of bonds | After | Num. of bonds |
| **Binding region** | Highly: 10.4%<br>Minimally: 28.5% | 298 | Highly: 5.6%<br>Minimally: 30.1% | 395 |
| **Dimer interface** | Highly: 13.7%<br>Minimally: 29.8% | 637 | Highly: 15.4%<br>Minimally: 25.4% | 625 |
|  | CPSF6 binding |  |  |  |
|  | Before | Num. of bonds | After | Num. of bonds |
| **Binding region** | Highly: 11.0%<br>Minimally: 30.1% | 4865 | Highly: 6.6%<br>Minimally: 34.4% | 5560 |
| **Dimer interface** | Highly: 14.7%<br>Minimally: 29.9% | 1120 | Highly: 9.05%<br>Minimally: 29.2% | 1215 |

### Residue-wise Frustration across the Binding Ligands

In addition to interface-specific and binding site-specific measurements, residue-level frustration patterns were analysed across the binding ligands. The Protein Frustratometer reports the proportion of contacts classified into each frustration category (highly frustrated, neutral, and minimally frustrated) within a sphere of 5 Å around each residue^28,29^. Rather than using these categories, we computed the average frustration index of all contacts surrounding each residue. This measure provides greater sensitivity for detecting subtle differences between Apo and ligand-bound states, which did not produce large shifts in categorical proportions but still alter underlying interaction energetics. The resulting residue-wise profiles are shown in Supplementary Figs. 16c and 17c.

### Monomer Alignment for CypA-binding site identification and Mapping

To identify and map CypA-binding monomers (via the canonical binding site), we applied the Needleman-Wunsch algorithm^69^, a dynamic programming method for sequence alignment. It is used here to compare the CypA-binding monomer from the CypA-CA complex (PDB: 6Y9W, Fig. 7b) with the three potential CypA-binding monomers in the Apo CA complex (PDB: 6SKN, Fig. 7a). Structural similarity was evaluated by root mean square deviation (RMSD). The monomer with the lowest RMSD value (0.002 Å, compared with 0.779 Å and 0.952 Å for the other two candidates) was identified as the corresponding CypA-binding monomer and coloured in cyan, while the other two were coloured in lime and yellow, respectively.

The same alignment procedure was used to map CypA-binding monomers from the CypA-CA complex (PDB: 6ZDJ, Supplementary Fig. 11e) onto the predicted binding monomers on the surface of the CA tubular structure (PDB: 6X63, Fig. 7c).

### Comparison of Local Curvature in Hexameric Surface lattices Using Covariance Matrices

To compare local curvature in different tubular hexameric surface lattices, we analysed data of CA hexamers from assemblies with distinct helical symmetries.

We first defined six triangles for each hexamer, as illustrated in Supplementary Fig. 12a (PDB: 4XFX). The centroid of each hexagon, determined from the N195 Cα atomic positions of the six CA monomers, was used as the central vertex. This point was then connected to each of its six outer vertices to form six triangles. In a planar hexamer without internal curvature (PDB: 4XFX), all six triangles are coplanar. By contrast, for curved hexamers (PDB: 6SKN and 6X63), they exhibit angular deviations, reflecting internal bending of the hexamer. In Supplementary Fig. 12b (PDB: 6SKN), six triangles are constructed for each hexamer and colour-coded according to the three monomers at the local threefold axis (as in Fig. 7a): hexagon 1 (yellow), hexagon 2 (cyan), and hexagon 3 (lime).

As illustrated in Supplementary Fig. 12b, here we define three hexamer array orientations^25^ as:

- ori1, the most curved direction, based on the hexamer arrangement composed of hexagons 1 and 2;
- ori2, the intermediate curvature direction, based on hexagons 1 and 3;
- ori3, the least curved direction, based on hexagons 2 and 3. Experimentally determined and theory predicted CypA-binding sites are both located at the dimer interface between hexagons 2 and 3.

As shown in the top row of Supplementary Fig. 13, in each orientation, we labelled triangles (tri1-tri6) for each orientation, starting from the dimer interface and ordered counterclockwise. To quantify curvature patterns, we computed 6 × 6 covariance matrices for each orientation, with entries corresponding to the relative angles between the normal vectors of the triangles in the adjacent hexamers. Analyses were performed on tubular assemblies with helical symmetries (−13, 8) (PDB: 6SKN; Supplementary Fig. 13, middle row) and (−12, 11) (PDB: 6X63; Supplementary Fig. 13, bottom row). Colour depth of each entry in the covariance matrix encodes the size of the angle.

The covariance matrices were approximately symmetric along the diagonal, implying a degree of symmetry along the local 2-fold axis between the hexamers. Deviations from perfect symmetry indicate relative distortions from perfect rotational symmetry. Averages over all matrix entries were taken as a proxy for the absolute curvature for each lattice orientation. In the (−13, 8) symmetry, two orientations exhibited higher curvatures (29.5° and 25.4°) and one lower curvature (5.9°), whereas in the (−12, 11) symmetry, one high (31.4°) and two lower curvatures (18.4° and 13.4°) were observed. Despite differences in absolute curvature values between these symmetries, the relative curvature pattern – meaning the locations of high and low values in the matrix – was preserved. This suggests that the relative geometric organisation revealed by this provides refine information superseding absolute curvature values.

Interestingly, the three relative orientations (ori1-ori3) correspond to the type 1-3 configurations defined by the geometric index (Supplementary Fig. 13, top row), demonstrating that the index reliably captures intrinsic local geometric organisation across different helical symmetries.

## Acknowledgments

The authors acknowledge funding from the Wellcome Trust 224509/A/21/Z (R.T.) and Exscientia (now Recursion; W.L.), and the US National Institutes of Health award R01AI178486 and U54AI170791 (J.R.P.). This work used the Extreme Science and Engineering Discovery Environment, which is supported by the National Science Foundation (Grant ACI-1548562). This work used ACCESS Delta and Stampede3 at the National Center for Super Computing Applications in Illinois and Texas Advanced Computing Center, respectively, through allocation MCB170096. Graphic visualisation was performed with UCSF ChimeraX^70^, developed by the Resource for Biocomputing, Visualization, and Informatics at the University of California, San Francisco, with support from National Institutes of Health R01-GM129325 and the Office of Cyber Infrastructure and Computational Biology, National Institute of Allergy and Infectious Diseases.

## Author Contributions Statement

W.L. & R.T. designed the study. W.L. developed the geometric index framework, lattice representations, and molecular frustration computations; performed the geometric and cofactor binding analyses; and created the figures. C.A.P., J.S.R., & J.R.P. carried out the carbon fullerene structure generation and corresponding energy computations. W.L. & R.T. interpreted the results and wrote the manuscript with input from all authors.

## Competing Interests Statement

The authors declare no competing interests.

## Data availability

The structural data generated in this study, together with the processed results, are available at Zenodo (carbon fullerene generation <u>DOI: 10.5281/zenodo.17654422</u> and others <u>DOI:</u> <u>10.5281/zenodo.17700975</u>). Other atomic structures used in this work were obtained from the Protein Data Bank under accession codes provided in the main text.

## Code availability

The source code used for carbon fullerene generation and optimisation is available at Zenodo (<u>DOI: 10.5281/zenodo.17654422</u>). All custom codes used for geometric index computation and analyses, lattice reconstruction, molecular frustration profiling, topography classification, local curvature quantification, and cofactor binding analyses are available at Zenodo (<u>DOI:</u> <u>10.5281/zenodo.17700975</u>).

## Notes

### Competing Interest Statement

The authors have declared no competing interest.

### Summary of Updates

Accepted for publication in Nature Communications. Revised the molecular frustration analysis to compare states before and after cofactor binding. Refined the colour scheme, formatting, and layout of selected figures and tables for improved clarity and consistency.

