## Supplementary Figures for "A geometric criterion links HIV-1 capsid topography to its biophysical properties and function"

### Supplementary Fig. 1

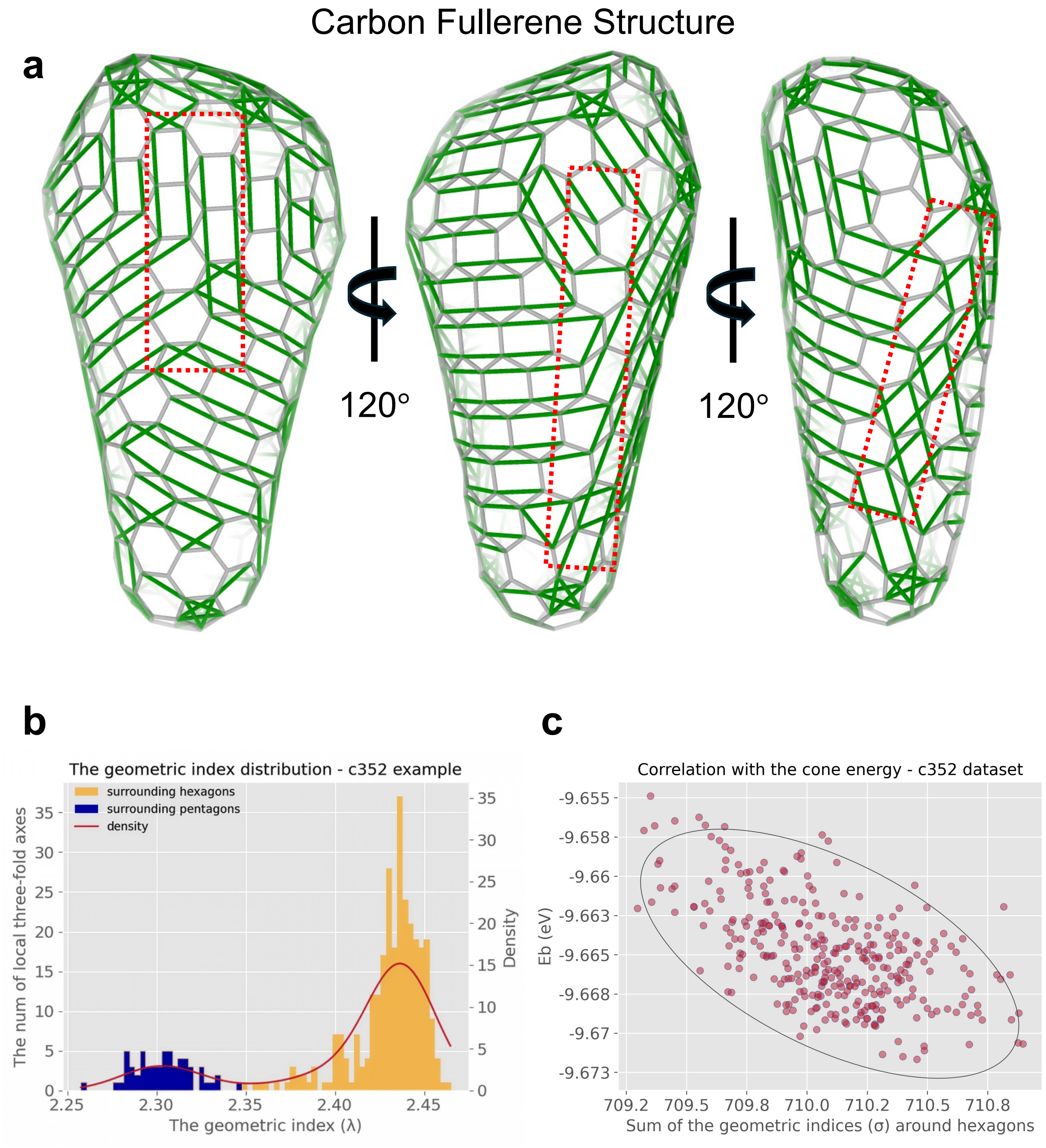


**Supplementary Fig. 1 | The geometric indices of a fullerene cone and their correlation with cone energy.** **a**, Graphic representation of the geometric indices (green lines) on the surface of a representative fullerene cone in the $C_{352}$ dataset. The geometric indices align to form partial rows, which form local sheets that are joined together at seam lines (see inside red dashed frames). **b**, The distribution of geometric indices of the cone in **a**. **c**, Cone energy correlates with the sum of the geometric indices surrounding hexagons for each cone in the $C_{352}$ dataset.

#### Supplementary Fig. 2

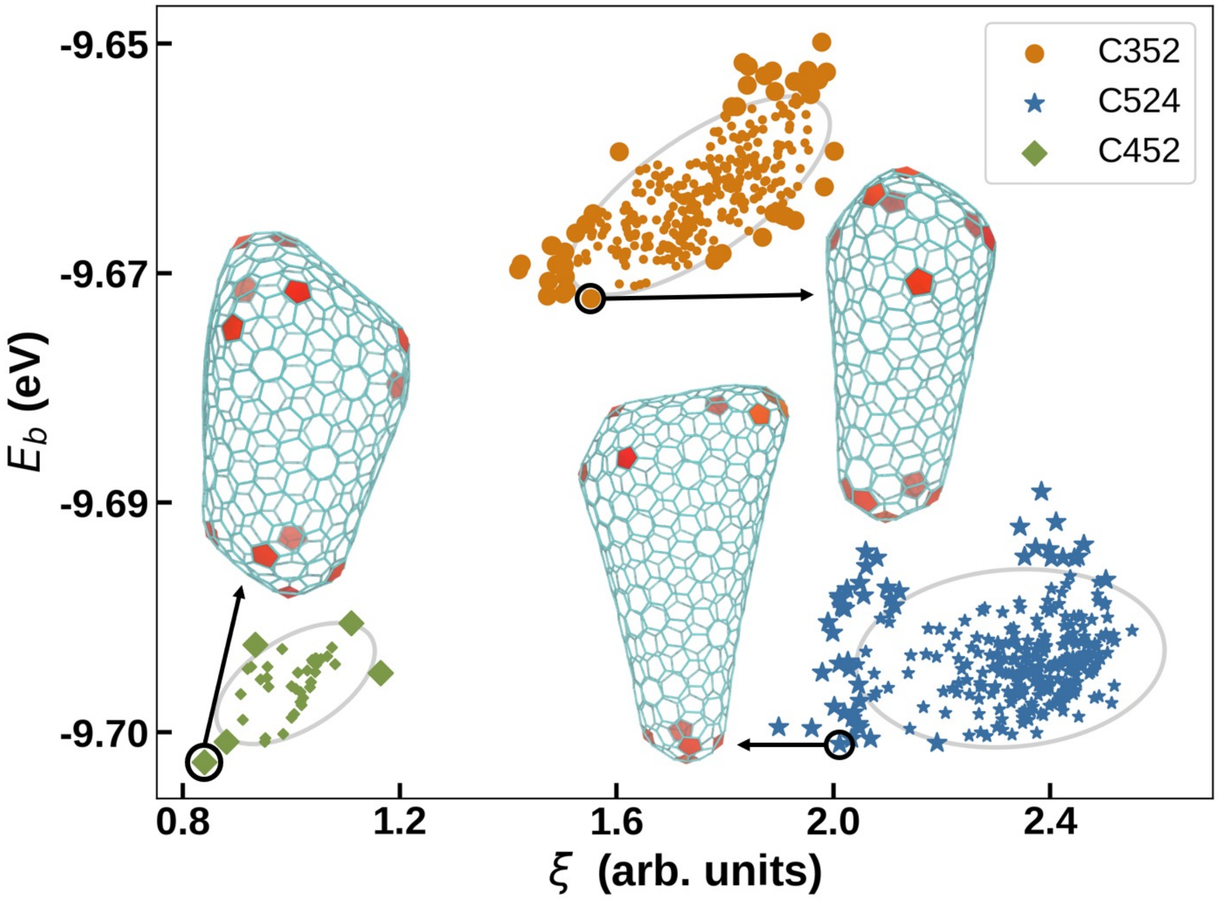


**Supplementary Fig. 2 | Correlation between the cone energy and a geometric parameter that describes the ratio between the width and height of the cone.** Each cone has exactly 12 pentamers, the grey circle indicates points which are 2 standard deviations within the mean (or ~95% of the data) of each set. The larger points are outside 2σ. The C-numbers in the legend describe how many carbon atoms are used to make each structure. The lowest energy structure for each set of fullerenes is highlighted by the black arrows.

### Supplementary Fig. 3

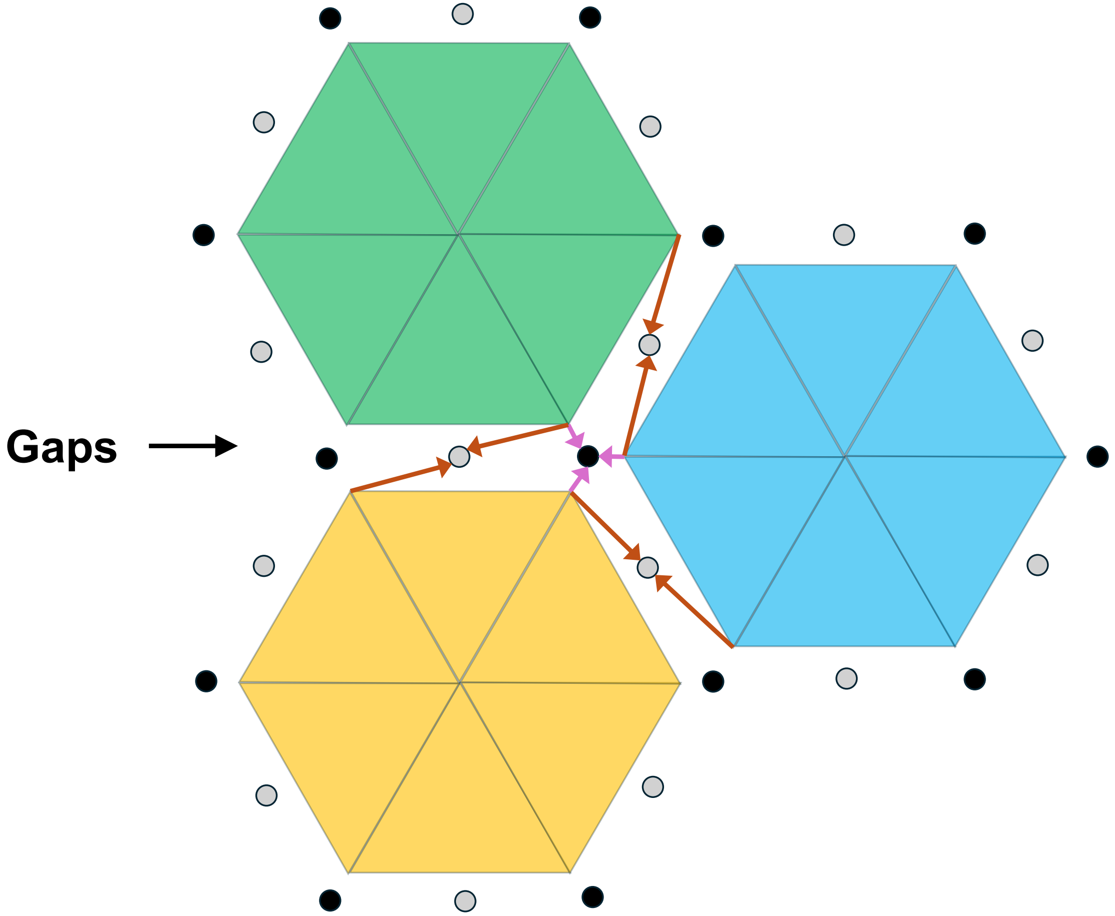


**Supplementary Fig. 3 | The construction of hexagonal and Kagome lattices based on three neighbouring hexagons.** Black dots correspond to the vertices of a hexagonal lattice. They are defined at the centroid of three vertices of the adjacent hexamers (magenta arrows). Grey dots mark the vertices of a Kagome lattice. They are defined as the midpoint between vertices from the two adjacent hexagons (brown arrows).

### Supplementary Fig. 4

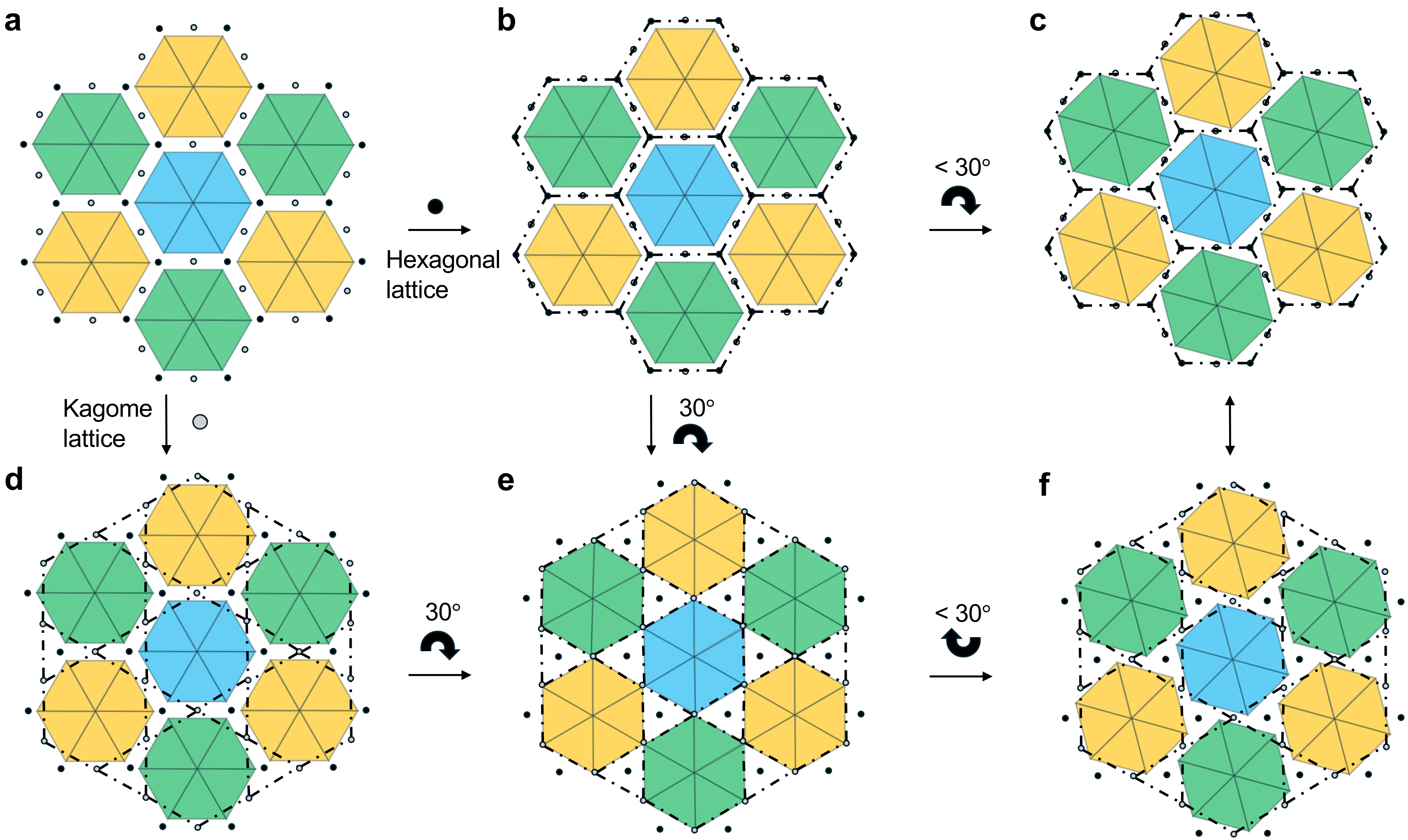


**Supplementary Fig. 4 | Fullerene, Kagome, and data-derive lattices in comparison. a,** A cluster of seven hexagons (coloured), with black and grey dots constructed based on the method in Supplementary Fig. 3-1. **b**, A hexagonal lattice derived from the black dots, where each enlarged hexagon encompasses the original (coloured) hexagon, covering the gaps. **c**, Coloured hexagons are rotated by < 30° about the centres of the original hexagons in **b**, mimicking the data-derived profile of CA hexamers, illustrating the difference to the hexagonal lattice. **d**, A Kagome lattice derived from the grey dots. **e**, Coloured hexagons are rotated by 30° about the centres of the original coloured hexagons in **d**, generating the characteristic triangular gaps of a Kagome lattice. **f**, Coloured hexagons are the same as in **c**, illustrating the differences between the Kagome lattice and the data-derived structure.

### Supplementary Fig. 5

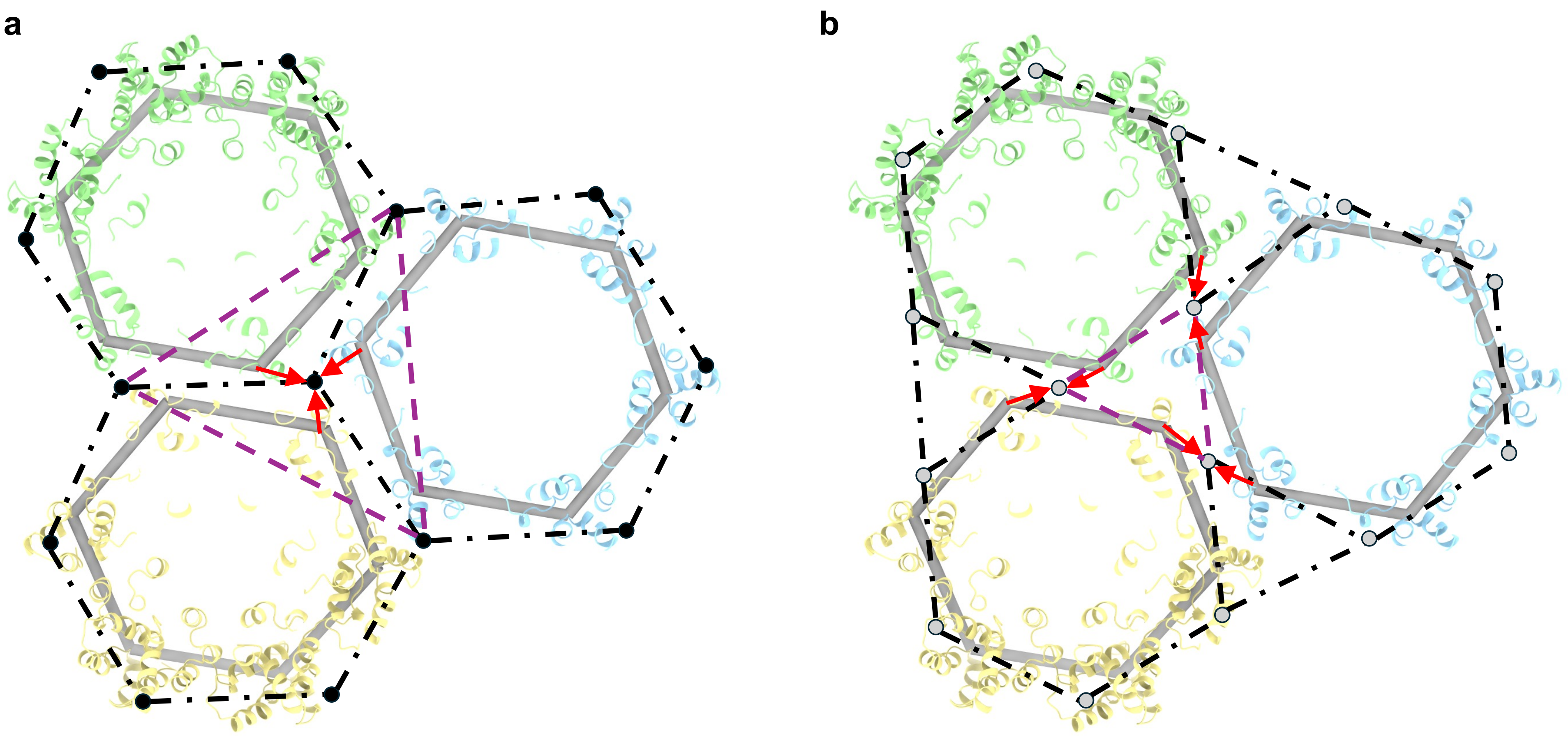


**Supplementary Fig. 5 | The fullerene, Kagome, and data-derived lattices in comparison.** Three hexamers (green, blue, and yellow) from vlp23 are shown, together with the data-derived model (light grey hexagons) constructed based on the N195 $C_\alpha$ atoms. Black and grey dots are derived based on the method in Supplementary Fig. 3-1. **a**, Comparison of the data-derived model with a fullerene lattice model, with the characteristic triangle of the fullerene lattice indicated by purple dashed lines. **b**, Comparison of the data-derived model with a Kagome lattice, with a characteristic triangle of the Kagome lattice shown as purple dashed lines.

### Supplementary Fig. 6

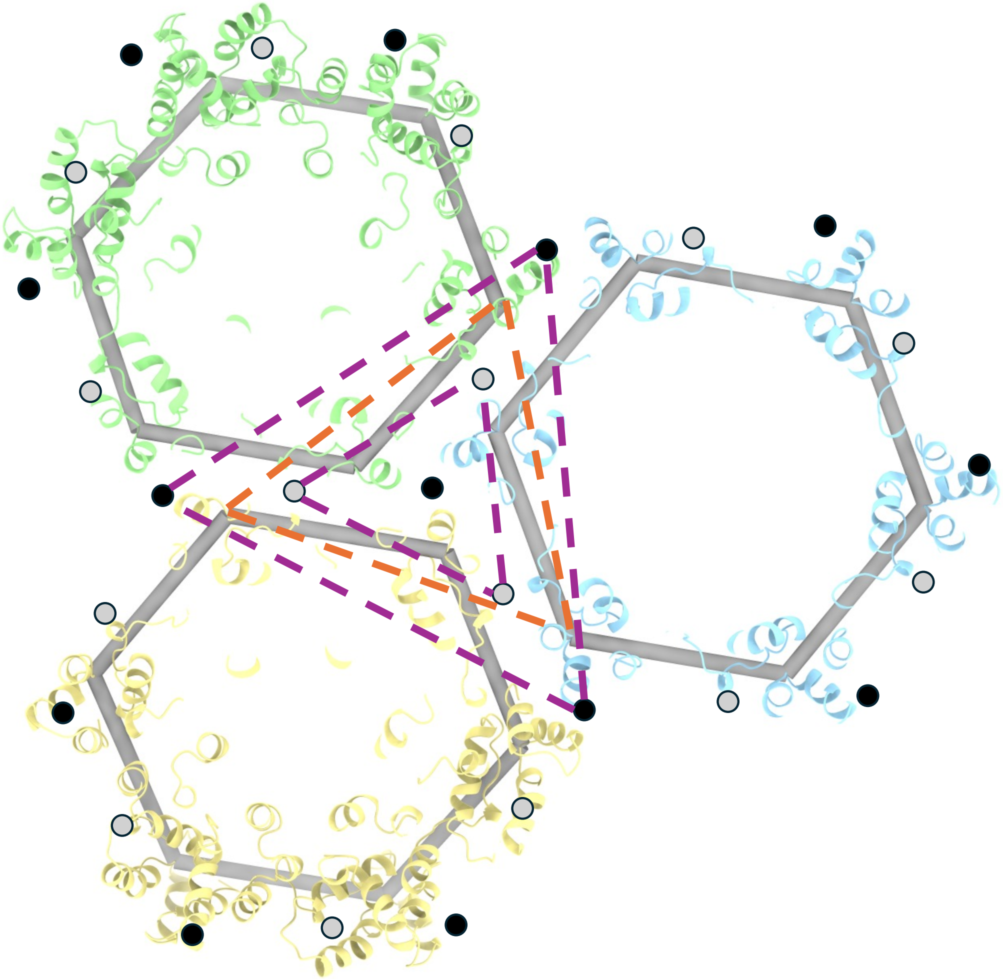

**Supplementary Fig. 6 | Comparison of the characteristic triangles of the fullerene, Kagome, and data-derived lattices.** Three hexamers (green, blue, and yellow) from vlp23 are shown in the context of the data-derived model (light grey hexagons) constructed from the N195 $C_\alpha$ atoms. Black and grey dots are constructed based on the method in Supplementary Fig. 3-1. Characteristic triangles for the fullerene and Kagome lattice are shown in purple dashed lines, whilst that of the data-derived model is shown in orange.

### Supplementary Fig. 7

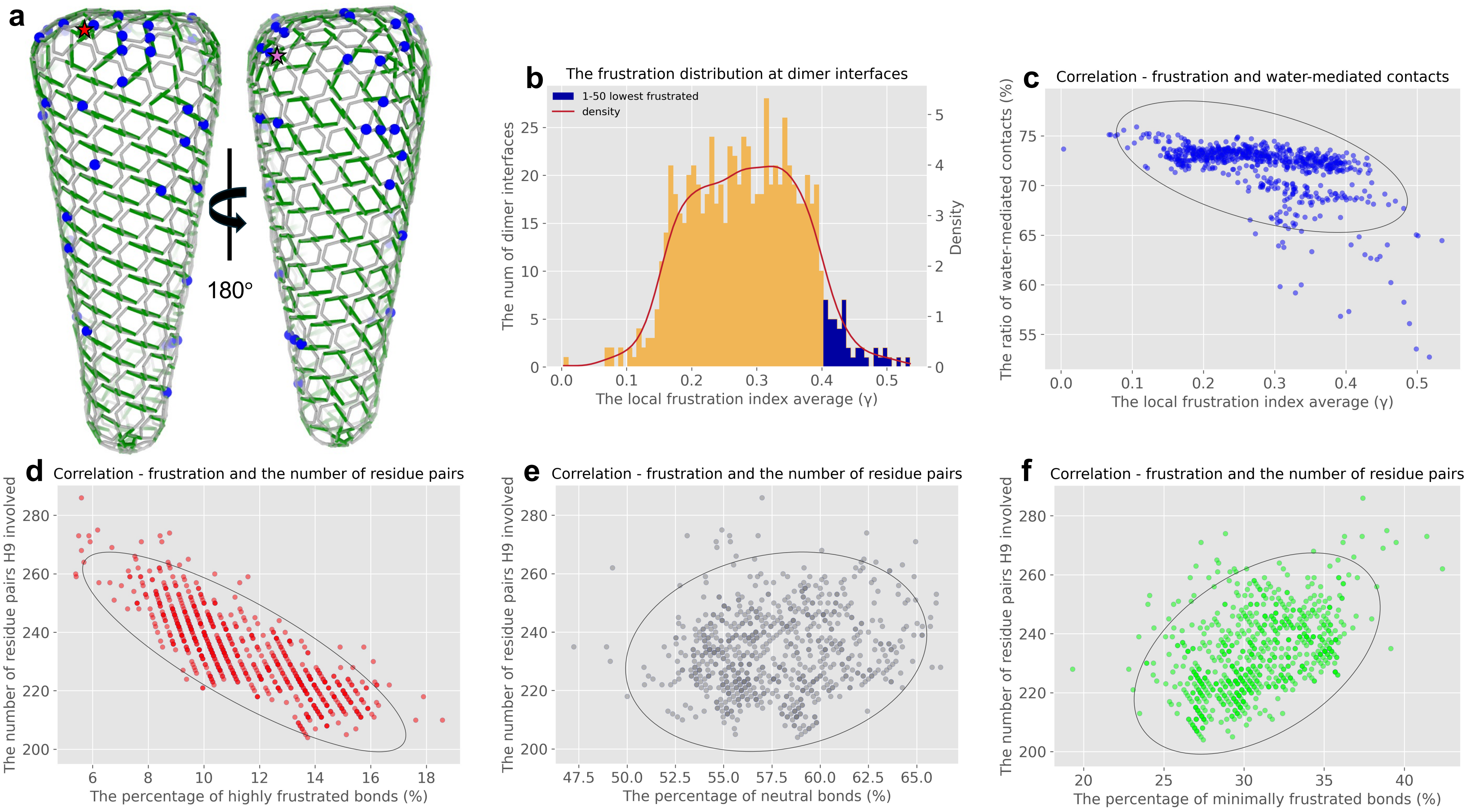


**Supplementary Fig. 7 | Distribution of minimally frustrated dimer interfaces and additional correlation analyses in *vlp23*. a**, The data-derived geometric model with the geometric indices in green. Blue spheres represent the 50 lowest frustrated dimer interface. Left: side view with the pentamer labelled by a red star, and Right: opposite side with the pentamer labelled by a light purple star, corresponding to the seam side. **b**, The distribution of the local frustration states across dimer interfaces in *vlp23,* colour-coordinated with **a**. **c**, Relationship between local frustration and the proportion of water-mediated contacts, showing a higher fraction of water-mediated interactions in more frustrated interfaces. **d-f**, Correlations between the number of residue pairs involved in H9 helix interactions and the percentage of highly frustrated (**d**), neutral (**e**), and minimally frustrated (**f**) bonds, showing that interfaces with more interactions exhibit fewer highly frustrated but more minimally frustrated contacts.

### Supplementary Fig. 8

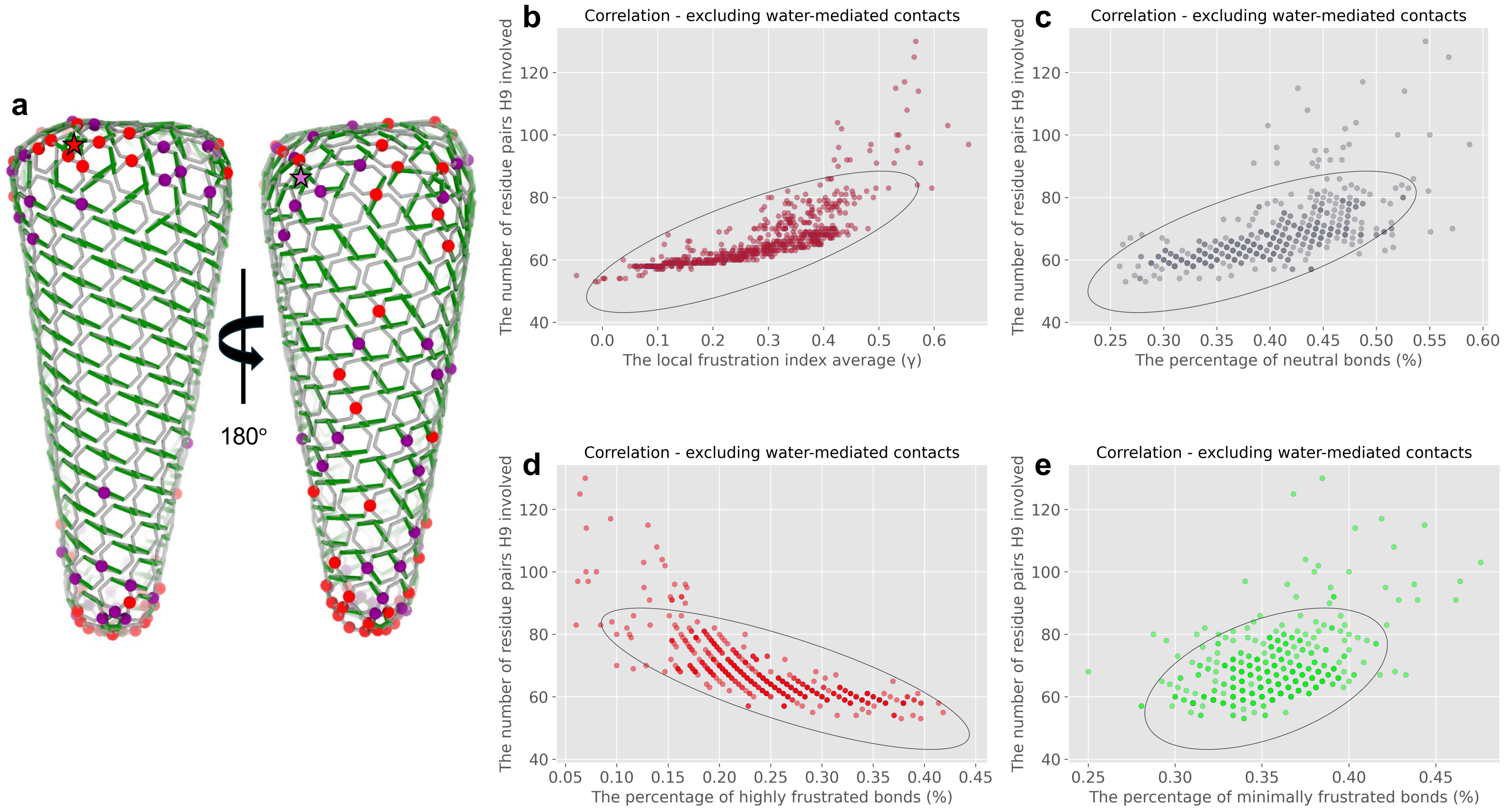


**Supplementary Fig. 8 | Molecular frustration at dimer interfaces excluding water-mediated contacts remains correlated with capsid topography given by the geometric index. a**, The data-derived geometric model of vlp23 with the geometric indices shown in green. Spheres indicate highly frustrated interfaces: red, top 50; purple, ranks 51-100. Left: side view with the pentamer labelled by a red star; Right: opposite side with the pentamer labelled by a light purple star, corresponding to the seam side. Highly frustrated interfaces in the bulk remain localised to the seam side even when water-mediated contacts are excluded. **b**, Relationship between local frustration and the number of residue pairs involved in H9 helix interactions, showing that interfaces with more contacts exhibit lower frustration even without water-mediated interactions. **c-e**, Correlations between the number of residue pairs involved in H9 helix interactions and the percentage of highly frustrated (**d**), neutral (**e**), and minimally frustrated (**f**) bonds, showing that interfaces with more interactions contain fewer highly frustrated but more neutral and minimally frustrated contacts.

### Supplementary Fig. 9

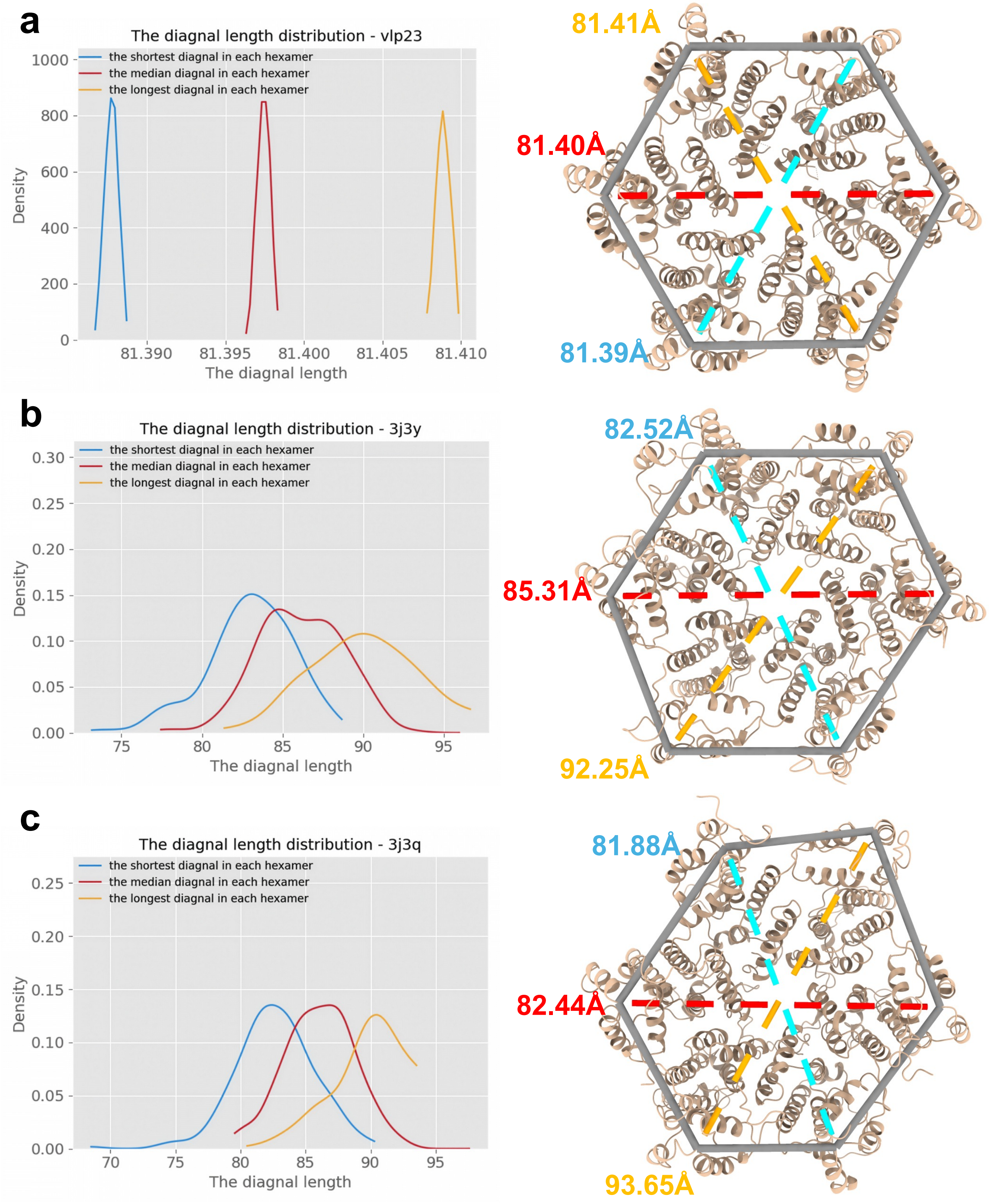


**Supplementary Fig. 9 | Variation in shape of the CA hexamers for different assembly scenarios. a**, Left: the length distribution of the diagonals of hexamers in the data-derived model based on vlp23. Right: a hexamer from vlp23 and its model representation (grey hexagon), with numbers denoting the lengths of the diagonals (dashed lines). **b**, Left: the length distribution of the diagonals of hexamers in the data-derived model based on 3J3Y. Right: a hexamer from 3J3Y and its model representation (grey hexagon), with numbers denoting the diagonal lengths (dashed lines). **c**, The corresponding results for 3J3Q.

### Supplementary Fig. 10

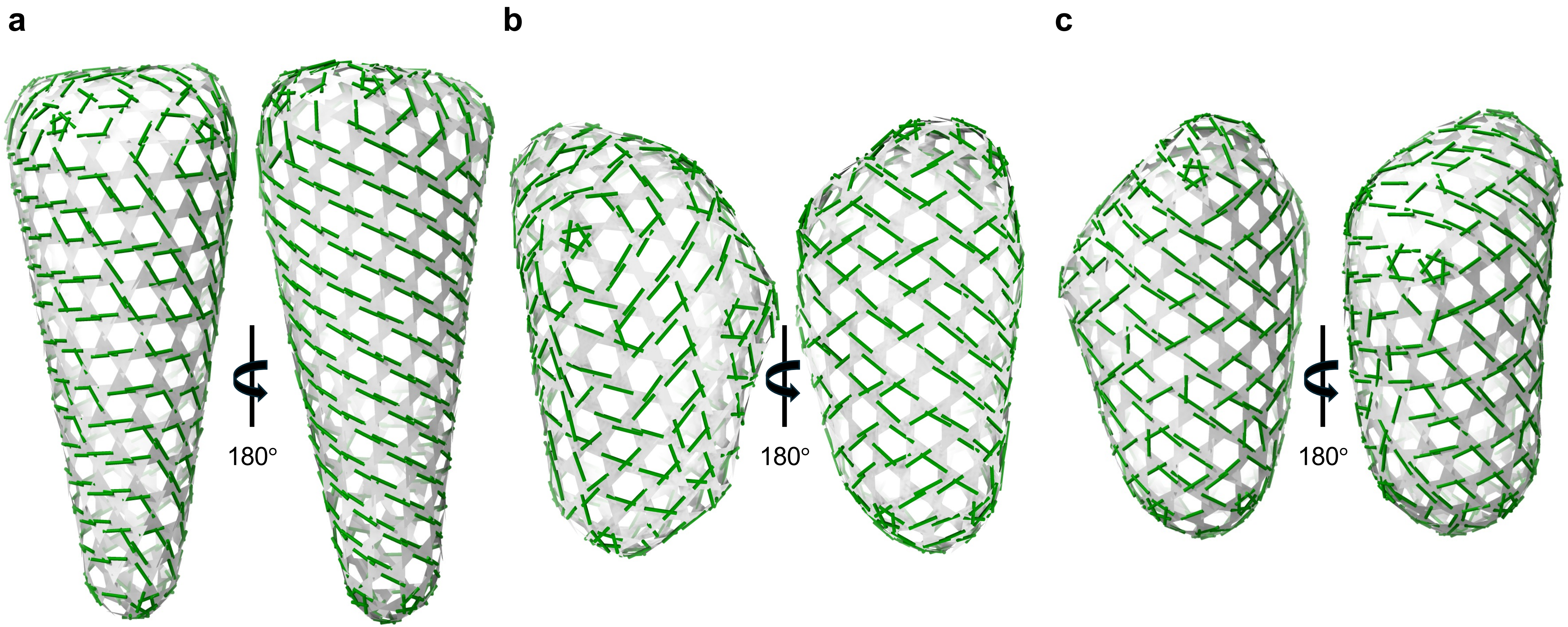


**Supplementary Fig. 10 | Geometric indices for different CA core morphologies. a**, Capsid structure of vlp23 showing the characteristic triangles (grey) of the data-derived model, with geometric indices superimposed as green lines. Capsid structures of **b**, 3J3Y and **c**, 3J3Q constructed along similar lines.

### Supplementary Fig. 11

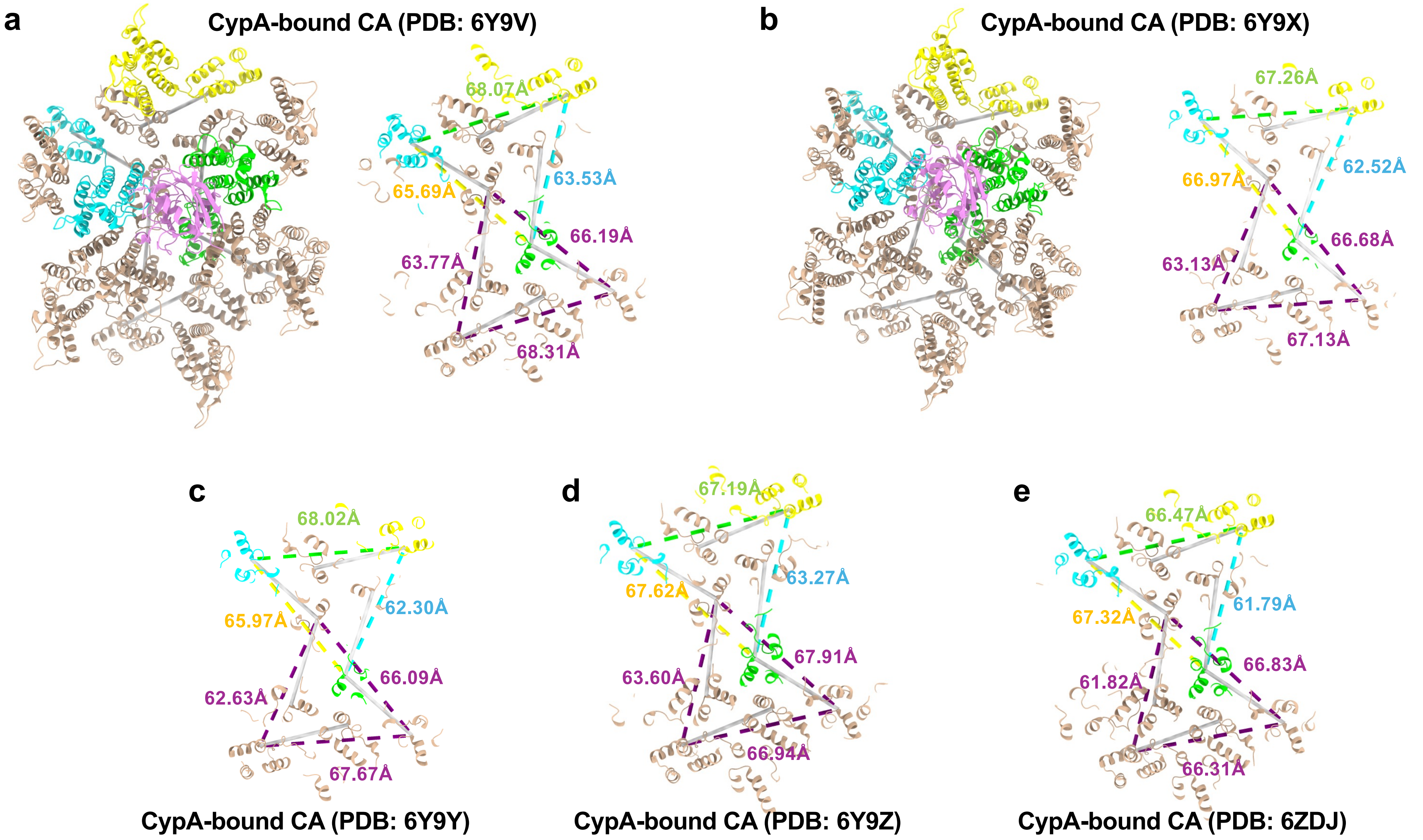


**Supplementary Fig. 11 | CypA-CA complexes in tubular structures with different helical symmetries.** Each of the complexes has two characteristic triangles (dashed lines) at two neighbouring local three-fold axes. The monomers coloured in cyan are the CypA-binding monomers via the canonical CypA-binding loops, with numbers denoting the lengths of the corresponding dashed lines. **a**, Helical symmetry (-8, 13); PDB: 6Y9V. **b**, Helical symmetry (-13, 7); PDB: 6Y9X. **c**, Helical symmetry (-7, 13); PDB: 6Y9Y. **d**, Helical symmetry (-13, 9); PDB: 6Y9Z. **e**, Helical symmetry (-13, 10); PDB: 6ZDJ.

### Supplementary Fig. 12

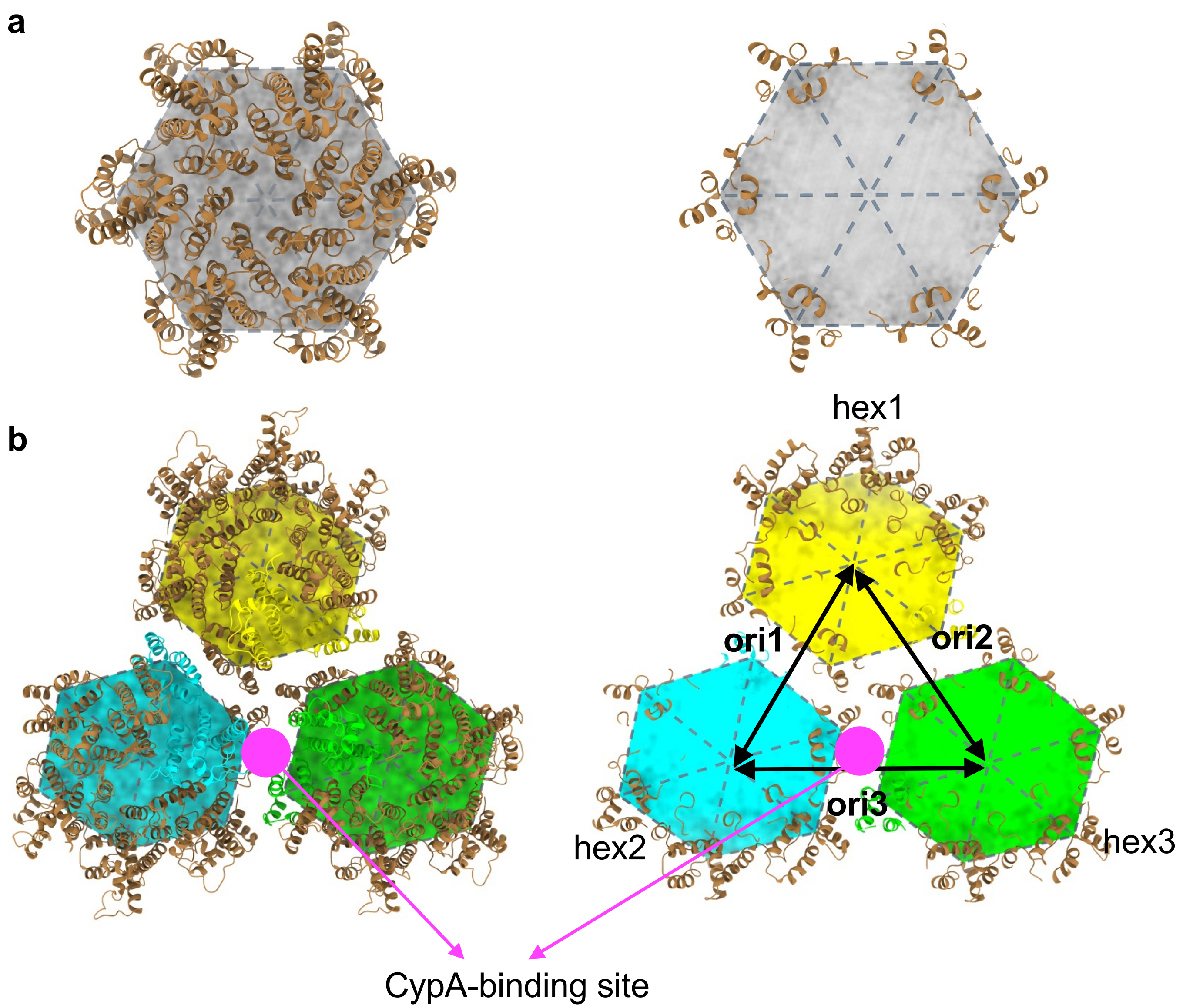


**Supplementary Fig. 12 | Schematics illustrating hexamer arrangements for curvature analysis. a**, Each hexamer is split into six internal triangles (grey) (PDB: 4XFX). The central vertex is defined as the centroid of the six outer vertices of the hexagon, and is connected to them (grey dashed lines) to form six triangles. In this flat reference model, all triangles lie in the same plane. **b**, Schematic of the same construction applied to three curved CA hexamers (PDB: 6SKN). Triangles are colour-coded to match hexagon 1 (yellow), hexagon 2 (cyan), and hexagon 3 (lime). Three hexamer array orientations are defined by connecting hexagonal midpoints as shown: ori1 (yellow-cyan, most curved), ori2 (yellow-lime, intermediate curvature), and ori3 (cyan-lime, least curved). The experimental and predicted CypA-binding sites are located at the dimer interface between hexagon 2 and 3 (ori3).

### Supplementary Fig. 13

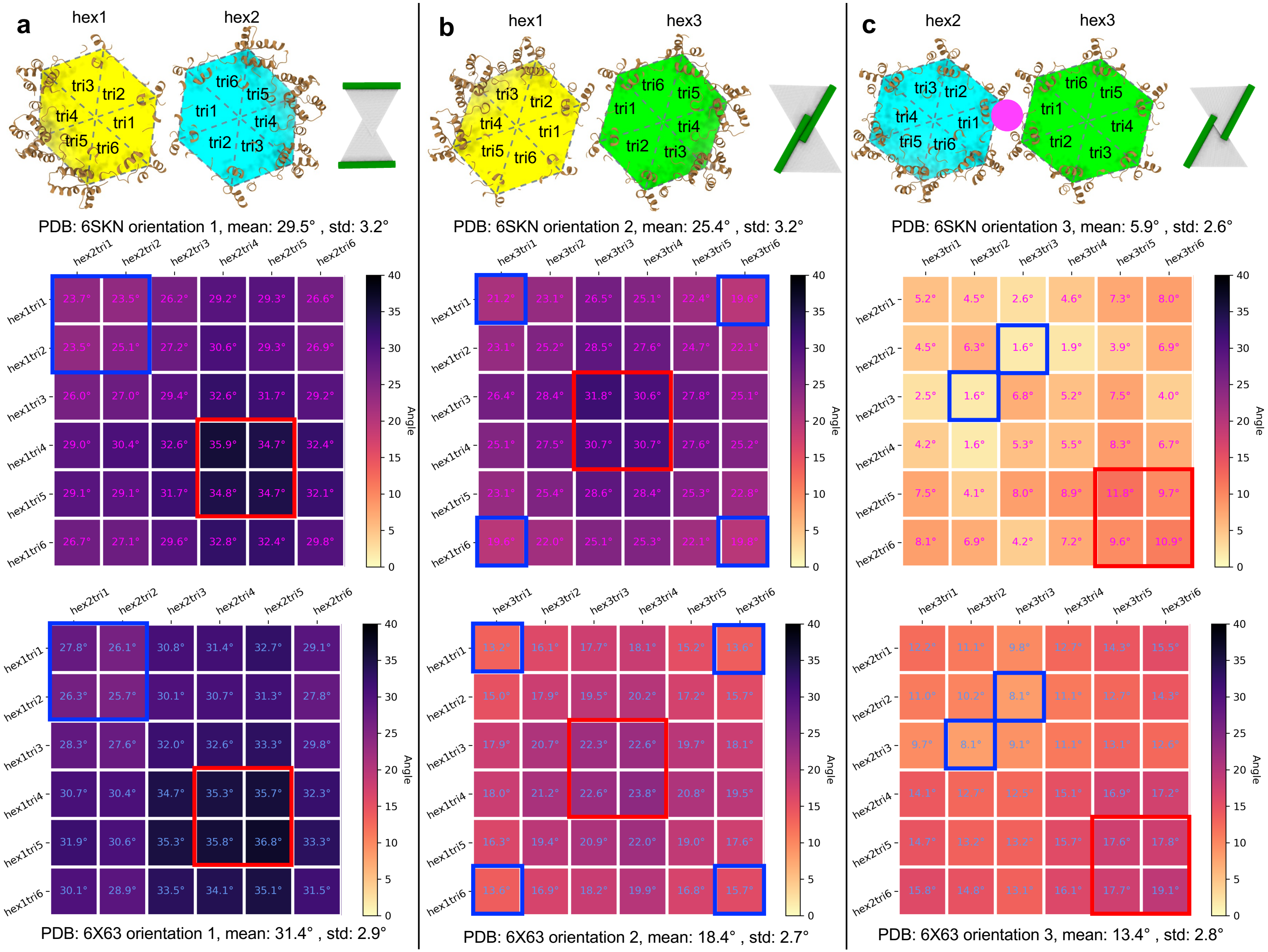


**Supplementary Fig. 13 | Covariance matrix comparing curvature across different tubular assemblies. a-c**, Top row: Illustation of hexamer arrangements along the three curvature orientations, from left to right: ori1-ori3, corresponding to type 1-3 configurations defined by the geometric index, respectively. Triangles are labelled (1-6) for each orientation, starting from the dimer interface and ordered counterclockwise (in keeping with the locations of N- and C-termini in the monomer). Middle row: Covariance matrices for each of the three orientations ori1-ori3, from left to right, in the CA tubular assembly with helical symmetry (-13, 8; PDB: 6SKN). Bottom row: Covariance matrices for helical symmetry (-12, 11; PDB: 6X63). Each $6 \times 6$ matrix represents angular relationships between the corresponding triangles in the two adjacent hexamers shown in **a**; colour depth encodes the size of the angle. Red frames highlight the largest angles (highest curvature), and blue frames the smallest (lowest curvature). Despite differences in absolute curvature between symmetries, the relative curvature patterns – high, mediate, and low – remain conserved.

#### Supplementary Fig. 14

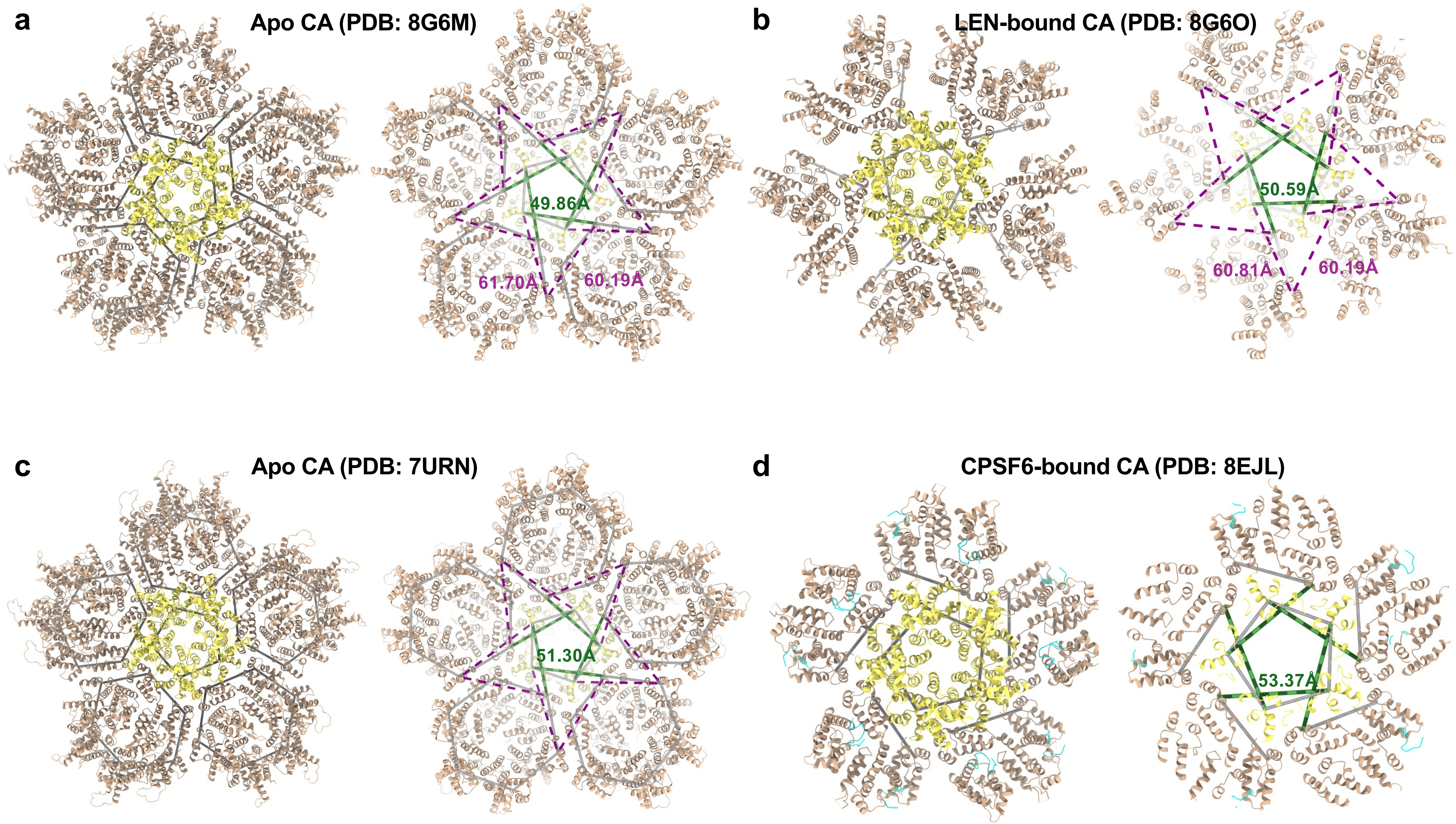


**Supplementary Fig. 14 | Structural implications of LEN and CPSF6 binding revealed by the geometric criterion.** The central pentamers (yellow) and surrounding hexamers (brown) are shown in the context of the data-derived model (grey lines). Geometric indices are highlighted in green (length given as a numeric value), with the purple dashed lines indicating the other edges of characteristic triangles. **a**, The HIV-1 CA lattice bound to IP6 at pH 7.4 (PDB: 8G6M). **b**, The HIV-1 CA lattice bound to IP6 and Lenacapavir at pH 7.4 (PDB: 8G6O). **c**, The HIV-1 WT CA capsid (PDB: 7URN). **d**, The HIV-1 CA capsid in complex with the CPSF6-FG peptide (PDB: 8EJL) where CPSF6s are coloured in cyan.

#### Supplementary Fig. 15

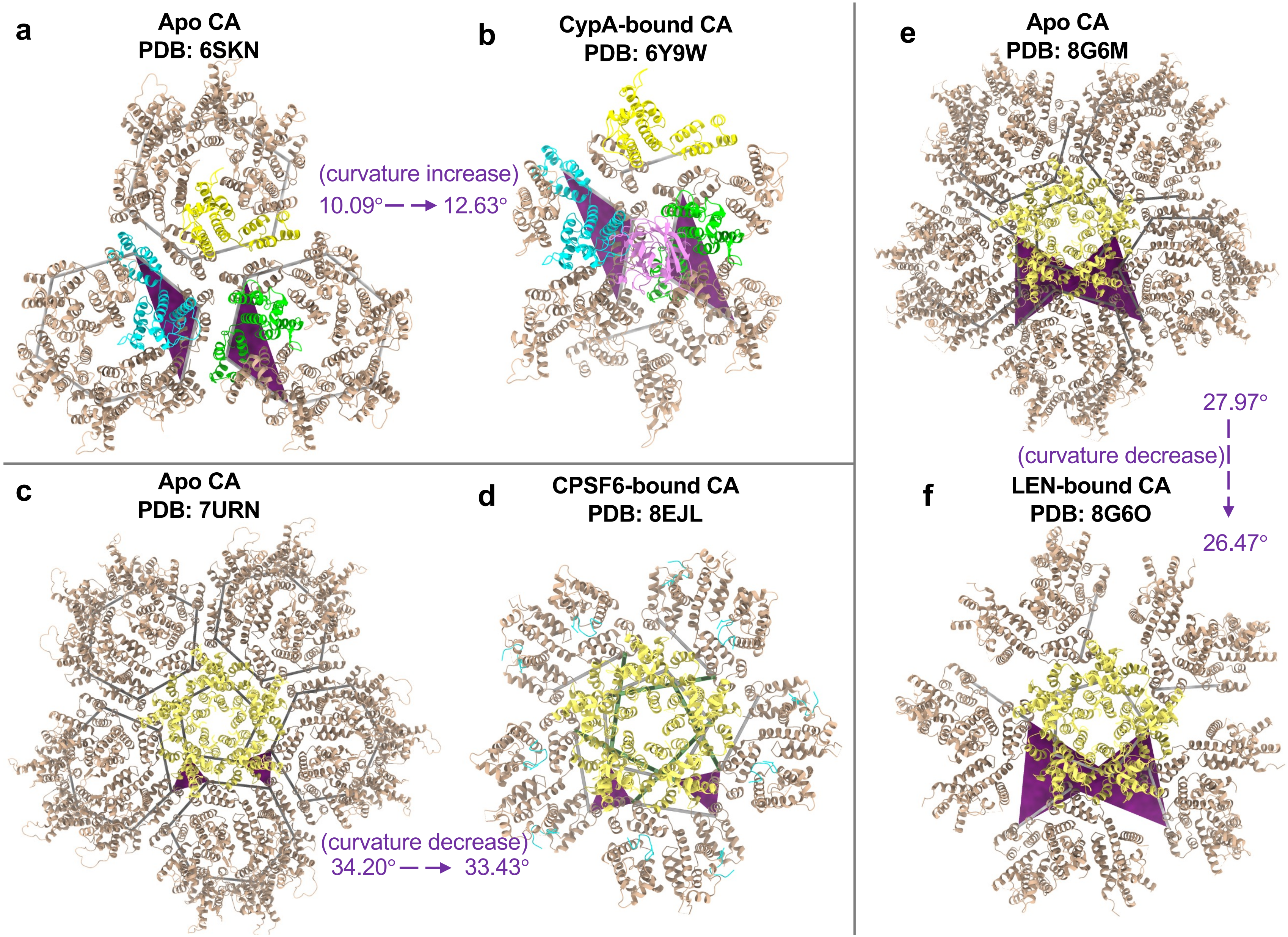


**Supplementary Fig. 15 | Structural effects of cofactor and small-molecule binding revealed by local curvature measurements.** Local curvature was quantified by defining two triangles (purple) in each structure and measuring the angle between their surface normal vectors. The specific triangles differ between cases due to structural constraints of the available PDB models before and after binding. CypA binding induces curvature enhancement, whereas CPSF6 and LEN binding produce local lattice flattening, consistent with the trends inferred from the geometric index. **a**, Apo CA complex prior to CypA binding (PDB: 6SKN). **b**, Corresponding CypA-CA complex (PDB: 6Y9W). **c**, Apo CA complex prior to CPSF6 binding (PDB: 7URN). **d**, Corresponding CPSF6-CA complex (PDB: 8EJL). **e**, Apo CA complex prior to LEN binding (PDB: 8G6M). **f**, Corresponding LEN-CA complex (PDB: 8G6O).

#### Supplementary Fig. 16

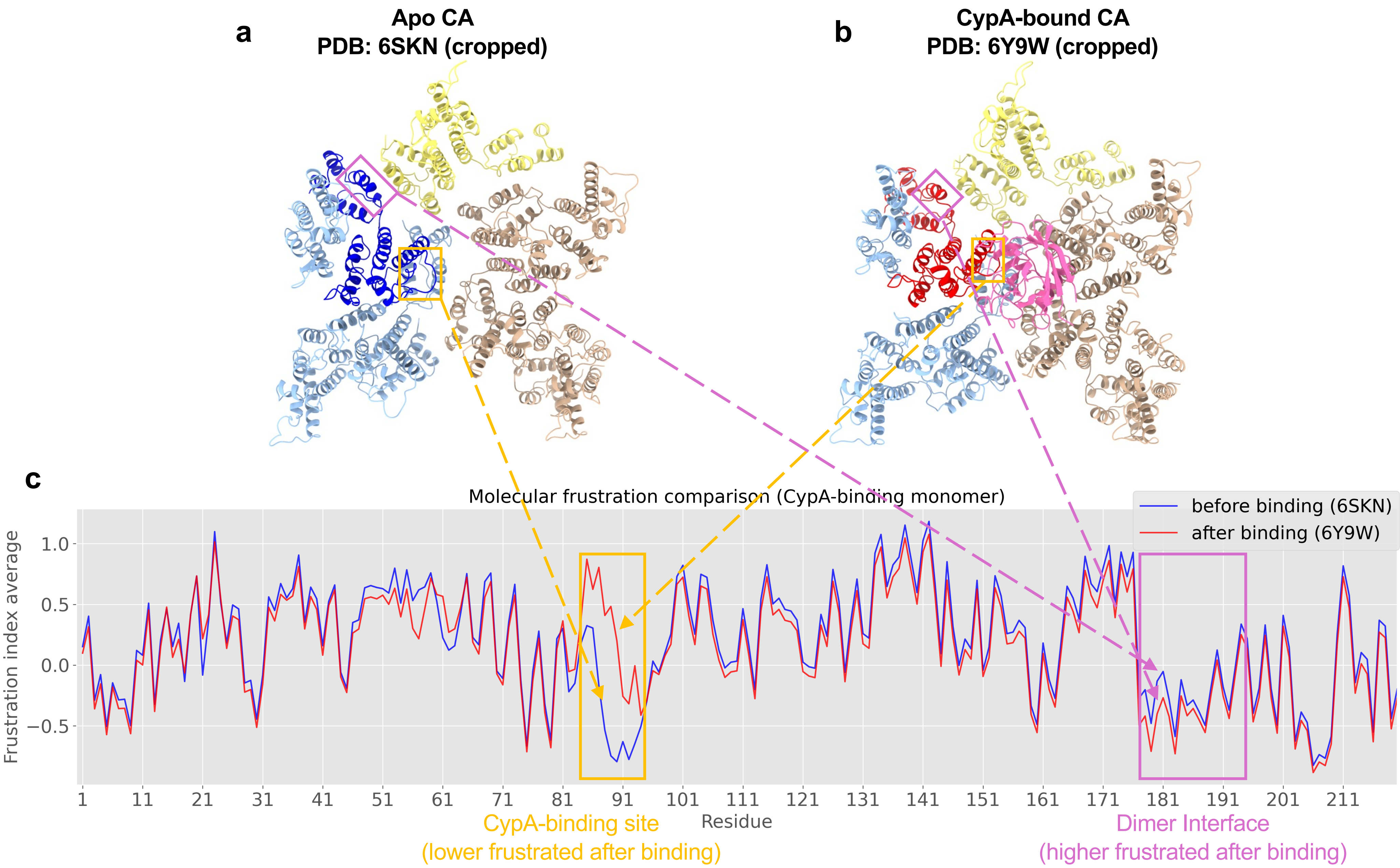


**Supplementary Fig. 16 | Molecular frustration comparison across the CypA-binding monomer.** **a**, Apo CA complex prior to CypA binding, with the CypA-binding monomer (via canonical binding site) highlighted in navy blue. **b**, CypA-CA complex, with the CypA-binding monomer highlighted in red. The CypA-binding site and dimer interface are indicated by orange and light purple frames, respectively. **c**, Residue-wise average molecular frustration index (residues 1-221) across the CypA-binding monomer before and after CypA binding.

#### Supplementary Fig. 17

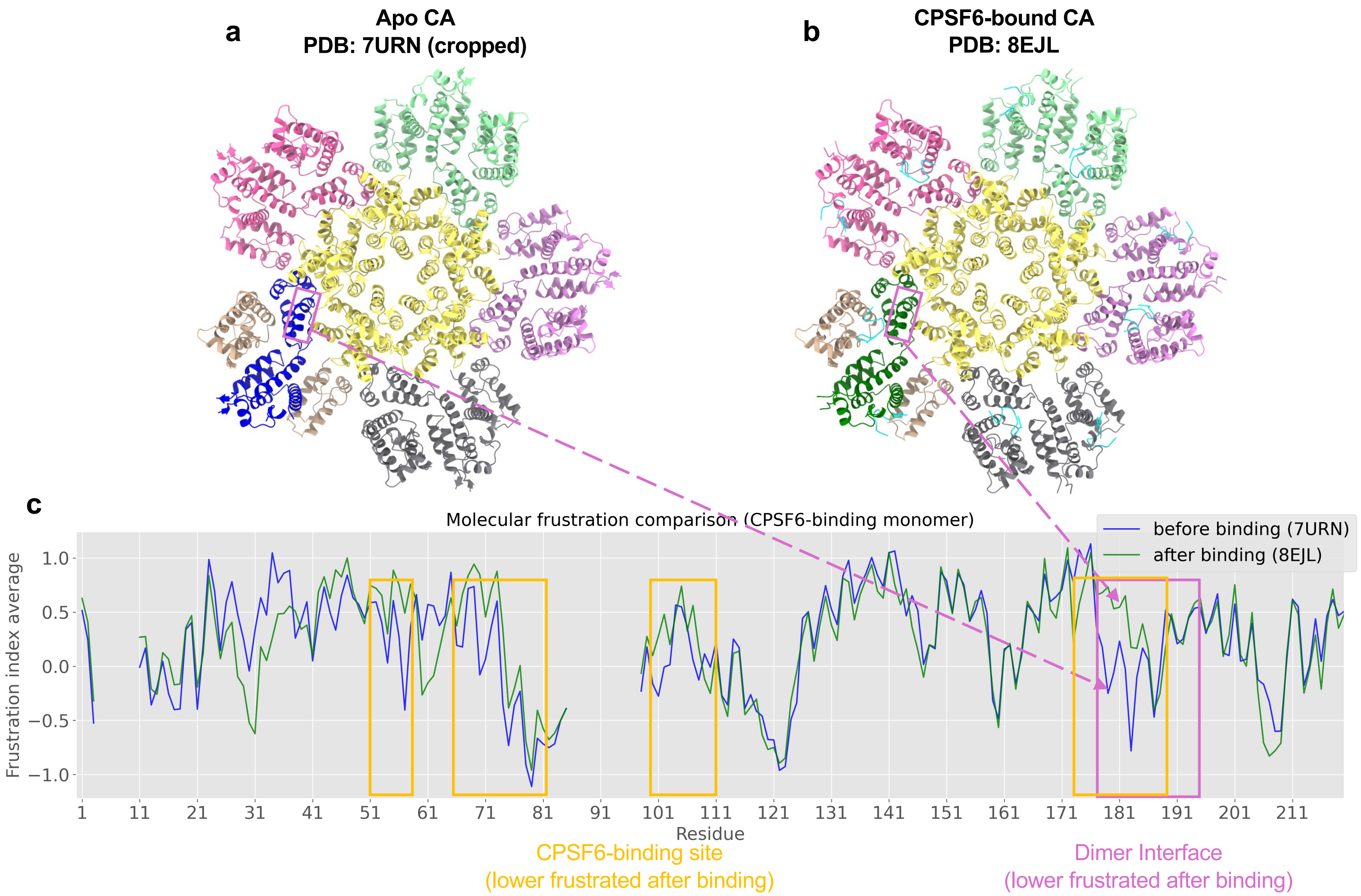


**Supplementary Fig. 17 | Molecular frustration comparison across the CPSF6-binding monomer. a**, Apo CA complex prior to CPSF6 binding, with one CPSF6-binding monomer highlighted in navy blue. **b**, CypA-CA complex, with one CPSF6-binding monomer highlighted in green. The CPSF6-binding site and dimer interface are indicated by orange and light purple frames, respectively. **c**, Residue-wise average molecular frustration index (residues 1-221; regions with missing residues omitted) across the CPSF6-binding monomer before and after CPSF6 binding.
