## Supplementary material for "A geometric criterion links HIV-1 capsid topography to its biophysical properties and function": Source Data

| Residues interacting with cofactors | | | | | |
|---|---|---|---|---|---|
| **CypA-bound complex (PDB: 6Y9W)** | | | **CPSF6-bound complex (PDB: 8EJL)** | | |
| Chain e | Chain K | Chain b | Chain L | Chain M | Chain N |
| 94 | 83 | 85 | 34 | 49 | 169 |
| 95 | 84 | 86 | 37 | 50 | 172 |
| 96 | 95 | 87 | 38 | 52 | 173 |
| 97 | 98 | 88 | 49 | 53 | 175 |
| 108 | 99 | 89 | 50 | 54 | 179 |
| 110 | 117 | 90 | 52 | 55 | 182 |
| 112 | 121 | 91 | 53 | 56 | 183 |
| 113 | 122 | 92 | 54 | 57 | 185 |
| | 123 | 93 | 55 | 63 | 186 |
| | 124 | 97 | 56 | 66 | 187 |
| | 125 | 100 | 57 | 67 | 211 |
| | 126 | 108 | 63 | 69 | |
| | 128 | | 66 | 70 | |
| | 129 | | 67 | 71 | |
| | | | 69 | 73 | |
| | | | 70 | 74 | |
| | | | 71 | 76 | |
| | | | 73 | 77 | |
| | | | 74 | 78 | |
| | | | 76 | 81 | |
| | | | 77 | 100 | |
| | | | 78 | 101 | |
| | | | 81 | 102 | |
| | | | 100 | 103 | |
| | | | 101 | 104 | |
| | | | 102 | 105 | |
| | | | 103 | 106 | |
| | | | 104 | 107 | |
| | | | 105 | 108 | |
| | | | 106 | 109 | |
| | | | 107 | | |
| | | | 108 | | |
| | | | 109 | | |
| | | | 169 | | |
| | | | 172 | | |
| | | | 173 | | |
| | | | 179 | | |
| | | | 180 | | |
| | | | 182 | | |
| | | | 183 | | |
| | | | 185 | | |
| | | | 186 | | |
| | | | 187 | | |
| | | | 211 | | |
